# A thalamic inhibitory circuit aligns sensory coding with learned value

**DOI:** 10.64898/2026.06.29.735190

**Authors:** Ricardo Paricio-Montesinos, Meike Knull, Amine Bahlouli, Jan Gründemann

## Abstract

Adaptive behavior requires sensory systems to prioritize cues that predict meaningful outcomes while suppressing irrelevant stimuli, but how relevance-based filtering is implemented along early sensory pathways remains unclear. Using deep brain two-photon imaging and causal circuit manipulations in mice performing an audiovisual detection task, we show that inhibition from thalamic reticular nucleus dynamically tunes sensory thalamus according to learned value. As animals learned stimulus-outcome associations, neurons in medial geniculate body developed biased responses favoring reward-predicting cues and suppressing non-rewarded stimuli. Silencing inhibitory input from thalamic reticular nucleus broadly disinhibited thalamic responses and abolished this value bias. Notably, stimulus identity decoding was unaffected by loss of inhibition. However, the geometry of MGB population activity was reorganized: coding axes rotated, action-related coding was strongly impaired, and thalamic representations became misaligned with learned behavioral readout. This altered population code impaired behavioral performance despite preserved sensory separability. Thus, inhibition of sensory thalamus by the reticular nucleus does not simply gate sensory throughput; it acts as a subcortical coding mechanism that aligns neuronal representations with learned value and behavioral goals, with implications for disorders of perception and cognition.

## Introduction

Animals learn which sensory cues predict reward, threat, or neutral outcomes, and selectively weigh those signals to guide behavior. Although experience- and relevance-based filtering is often attributed to cortical networks^1–7^, learning also reshapes neural representations already in sensory thalamus^1–4,8^. However, the circuit mechanisms that implement flexible, experience-dependent coding at thalamic sensory processing stages remain unknown.

Inhibition is a fundamental computational mechanism for adaptive learning^5,9^. In sensory thalamus, inhibitory control is mediated primarily by the thalamic reticular nucleus (TRN)^10,11^, which provides major inhibitory input to thalamic nuclei including the auditory thalamus (medial geniculate body, MGB).

TRN receives convergent projections from sensory^12^, executive^13,14^ and limbic regions^15^, and has long been proposed to regulate sensory selection in thalamic nuclei^16–18^. This positions TRN inputs as a candidate circuit for aligning thalamic representations with learned value, shaping population activity to guide outcome-dependent behavior. Here we test whether inhibitory input from TRN regulates learning-dependent plasticity in auditory thalamus and supports value-based sensory gating during behavior.

## Results

### Auditory thalamus activity is shaped by TRN inhibition in naïve animals

To understand how TRN regulates the coding of novel sensory cues in auditory thalamus of naïve animals, we imaged the activity of large populations of individual neurons in the MGB of awake, head-fixed mice using deep-brain two-photon calcium imaging, while silencing TRN chemogenetically. Since parvalbumin-expressing GABAergic neurons (PV^+^) constitute the majority of the anatomical TRN input to MGB (~88%)^19^, and represent the principal functional output to MGB^20^, we expressed the inhibitory DREADD hM4D(Gi) selectively in TRN of PV-Cre mice to silence TRN-mediated inhibition in MGB during passive stimulus exposure (Figure 1A). To simultaneously monitor MGB neuronal activity, we imaged GCaMP8m fluorescence changes of individual neurons through a GRIN lens above MGB (Figure 1A, S1). Postmortem anatomical tracing confirmed dense innervation of MGB by DREADD-expressing PV^+^ TRN axons (Figure 1B, S2).

**Figure 1.**
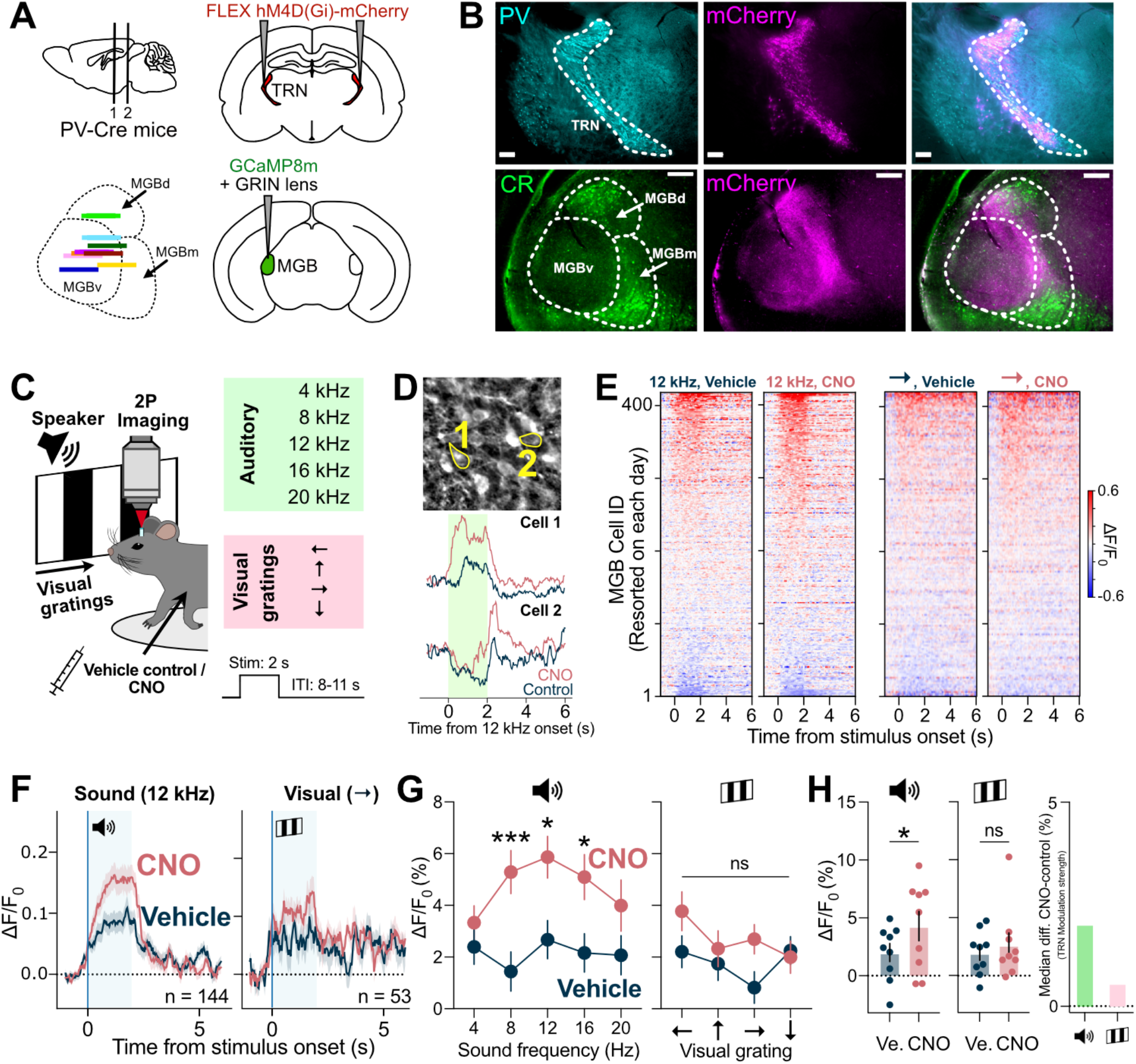
TRN suppresses responses to sound in naïve auditory thalamus. (**A**) Inhibitory DREADDs were expressed bilaterally in TRN of PV-Cre mice, and GCaMP8m in auditory thalamus (MGB). A GRIN lens was implanted above MGB (lens locations, bottom left). (**B**) Immunofluorescence confirms DREADD expression in PV+ TRN neurons (top) and dense mCherry-positive projections in MGB (bottom, calretinin-labeled). (**C**) Two-photon imaging of auditory thalamus neurons in head-fixed mice during presentation of auditory (4-20 kHz tones) and visual (drifting gratings) stimuli (2 s, interleaved; 8-11 s interval) following CNO or vehicle injection. (**D**) Individual neurons were tracked across sessions. (**E**) Example peri-stimulus time histogram (PSTH) heatmaps showing MGB responses to auditory (12 kHz) and visual (→ grating) stimuli under control and TRN blockade. Cells are sorted by stimulus-period response within each session. (**F**) Mean population responses to auditory and visual stimuli under control and TRN blockade. Included cells were those classified as responsive to sound (12 kHz) or visual (**→**) stimuli in either condition. (**G**) TRN blockade increased auditory responses (8, 12, 16 kHz) but did not significantly affect visual responses (two-way ANOVA, main effect of condition F(1,4902) = 26.54, p<0.0001, Bonferroni post hoc correction: 8 kHz p < 0.001, 12 kHz p = 0.0238, 16 kHz p = 0.0238; n = 416 cells). (**H**) Similar effects were observed when pooling all auditory and visual stimuli for each mouse (paired t test, p = 0.0150; n = 9 mice).

Mice were exposed to auditory (4, 8, 12, 16 and 20 kHz pure tones), visual (drifting gratings in four orientations) as well as multisensory stimuli (Figure 1C), given that sensory thalamus can respond to extramodal inputs^1,2,21^. Individual neurons were tracked across vehicle control and CNO injection sessions (Figure 1D).

Chemogenetic blockade of TRN increased sound-evoked responses in MGB neurons, while visual responses were comparatively less affected (Figure 1E-1H). In addition to increased response amplitudes, TRN silencing enhanced recruitment of neuronal responses for both auditory and visual stimuli (Figure S3A-S3B) and increased response gain among sound-responsive neurons (Figure S3C-S3D). Despite amplified multisensory responses (Figure S4A), the multisensory linearity index remained unchanged, indicating preserved sublinear integration (Figure S4B). Control mice expressing mCherry in TRN showed no differences between CNO and vehicle injection (Figure S2E).

Together, these results demonstrate that TRN suppresses auditory responses in naïve auditory thalamus while exerting comparatively weaker effects on visual inputs, establishing TRN as a modality-biased regulator of early thalamic stimulus processing.

### Upon learning, thalamic suppression becomes relevance-dependent

We next asked whether TRN modulation over MGB is static or whether it is dynamically shaped by learned stimulus relevance. To test this, we imaged MGB neurons during behavior in PV-Cre mice expressing hM4D(Gi) in TRN (see Figure 1) and trained on a Go/No-Go task. Sound-trained mice learned to lick for a 12-kHz tone predictive of reward and to withhold licking to a visual grating (Figure 2A-2C). Each trial consisted of a 2 s stimulus, a 2 s delay and a 1.5 s response period. Neuronal activity was recorded in expert mice during vehicle (control) and CNO (TRN blockade) sessions (Figure 2A). To compare TRN modulation across task contingencies, a separate cohort was trained on the reversed rules, with the visual cue predicting reward (visual-trained mice, Figure 2B).

**Figure 2.**
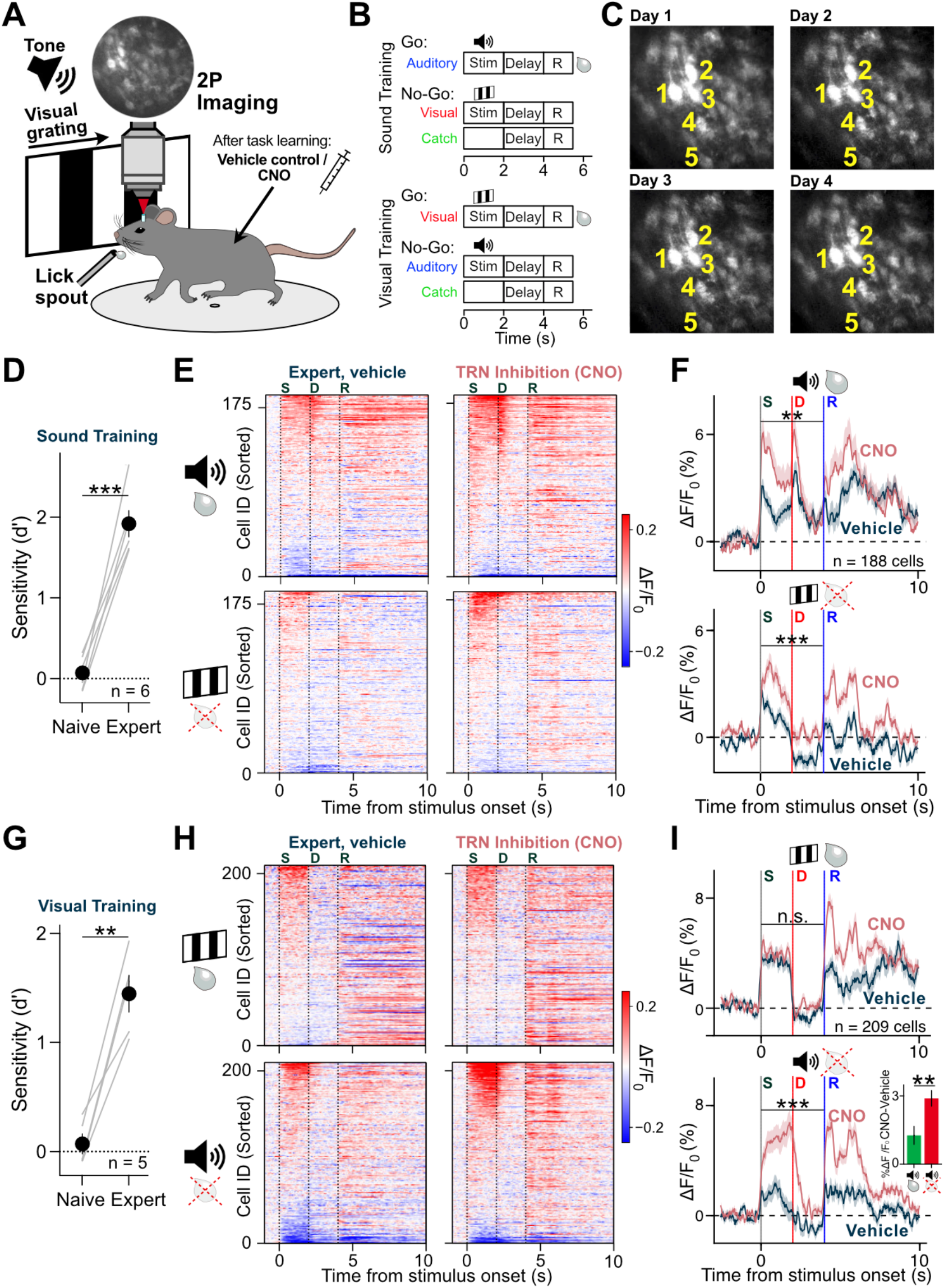
TRN suppression of MGB cross-modal responses becomes context-dependent upon learning. (**A**) Two-photon imaging of MGB neurons in head-fixed mice performing a Go/No-Go detection task. Mice reported trials by licking. At expert performance, animals were tested under CNO or vehicle. (**B**)Trial structure. Sound-trained mice reported auditory (12 kHz) trials and withheld licking to visual (**→** grating) and catch trials; visual-trained mice reported visual trials. Trials (interleaved) consisted of stimulus (2 s), delay (2 s) and response (1.5 s) epochs. Correct Go responses were rewarded. (**C**)Example field of view across four sessions (different days) showing stable cell tracking. (**D**) Sound-trained mice learned from naïve (d’ ~ 0) to expert performance (mean d’ = 1.92, paired t test p < 0.001; n = 6 mice). (**E**) PSTH heatmaps of MGB responses in sound-trained mice to auditory (12 kHz) and visual (→) trials under control and TRN blockade. Cells are sorted by stimulus-period response; bands at 4.5 s and 5.5 s reflect lick spout movement. (**F**) Mean population responses in sound-trained mice to sound (Go) and visual (No-Go) trials under control (black) and TRN blockade (red). TRN blockade increased responses (stimulus + delay) to both trial types (Mann-Whitney U: sound Go p = 0.006, visual No-Go p < 0.001; n = 188 cells, 5 mice). (**G**) Visual-trained mice learned from naïve (d’ ~ 0) to expert performance (mean d’ = 1.45, paired t test p = 0.0019; n = 5 mice). (**H**) MGB response PSTHs in visual-trained mice under control and TRN blockade. (**I**) Population responses in visual-trained mice. TRN blockade increased auditory (No-Go) but not visual (Go) responses (Mann-Whitney U, p < 0.001; n = 209 cells, 4 mice). Inset, comparison of TRN modulation of sound responses between sound-trained and visual-trained mice (p = 0.0091).

In expert sound-trained mice, TRN blockade enhanced MGB responses to both the auditory Go cue and the visual No-Go cue (Figure 2D-2F). In contrast, in expert visual-trained mice, TRN silencing selectively enhanced responses to the auditory No-Go cue, leaving visual Go responses largely unchanged (Figure 2G-2I). Notably, the magnitude of TRN-mediated auditory suppression was greater when sound served as the No-Go cue than when it predicted reward (Figure 2I). Control mice expressing mCherry in TRN showed no differences between CNO and vehicle sessions (Figure S2F).

These results indicate that TRN provides dynamic inhibition over auditory thalamus, which depends on learned context. Thalamic encoding of stimuli learned to be irrelevant (No-Go) is consistently suppressed by TRN in expert mice, regardless of sensory modality. However, for reward-predicting cues, TRN exerts a modality-specific suppression over auditory thalamus. Auditory Go cues are still suppressed (albeit with less strength than No-Go), but extramodal, visual Go cues are unaltered. Thus, TRN flexibly shapes thalamic coding upon learning to reflect task contingencies and behavioral demands.

### Inhibition from TRN regulates thalamic coding and learned behavior

Because TRN suppressed responses to familiar cues, we initially hypothesized that silencing TRN might facilitate detection and improve task performance. Instead, CNO injection caused a pronounced reduction in behavioral accuracy in both sound-trained (Figure 3A) and visual-trained mice (Figure 3B). The reduction in accuracy (measured as sensitivity, or d’) was driven by failures to report the Go cue rather than increased licking to the No-Go stimulus, indicating preserved task engagement (Figure 3A-3B). Control mice expressing mCherry in TRN showed no behavioral performance change upon CNO injection (Figure 3C).

**Figure 3.**
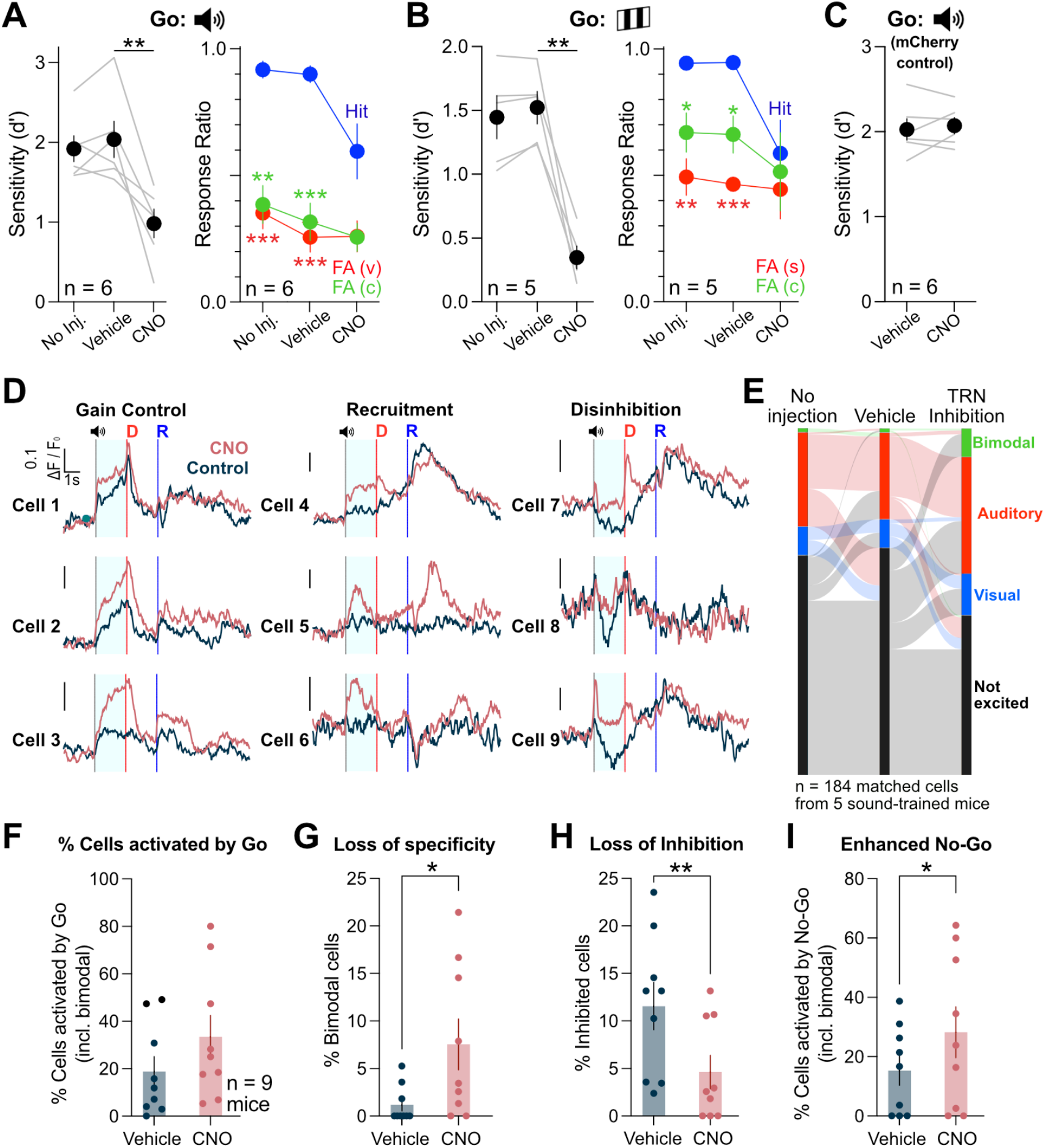
Inhibition from TRN guides learned behavior and single-cell coding in MGB. (**A**) Behavioral sensitivity (d’) in sound-trained expert mice. Vehicle had no effect, whereas CNO reduced performance (RM-ANOVA, main effect of treatment F(1.294, 6.471) = 20.16, p = 0.0026; Tukey post hoc: CNO vs vehicle p = 0.0152, vs no injection p = 0.0065, vehicle vs no injection p=0.6161; n = 6 mice). Right: hit and false-alarm rates differed significantly under no injection (all p ≤ 0.0011), but not under CNO. (**B**) Same for visual-trained mice (RM ANOVA, F(1.046, 4.183) = 39.26, p = 0.0028). CNO reduced performance (CNO vs vehicle p = 0.0044, vs no injection p = 0.01; n = 5 mice, Tukey post hoc test). Right, CNO reduced hit vs false-alarm separation (no injection p = 0.006 FA_(V)_, p = 0.047 FA_(C)_; vehicle p < 0.001 FA_(V)_, p = 0.035 FA_(C)_). (**C**) mCherry controls showed no effect of CNO. (**D**) Example neurons illustrating gain increase, response recruitment and disinhibition. (**E**) Sankey plot of response-class transitions across conditions. (**F**) Across all expert cells, CNO induced a non-significant trend toward increased Go-responsive cells. (**G**-**I**), TRN blockade increased bimodal neurons (Wilcoxon matched-pairs, p = 0.0156) and No-Go excited cells (p = 0.0156), and decreased stimulus-inhibited cells (p = 0.0078, n = 9 mice).

We next investigated single-cell changes in auditory thalamus in expert mice upon TRN inactivation. Blockade of TRN produced three prominent effects in MGB neurons: gain modulation (preserved tuning but with larger amplitudes), response recruitment (previously unresponsive neurons becoming active) and disinhibition (loss of stimulus-evoked suppression) (Figure 3D). Changes in responsiveness between control sessions were minor by comparison to TRN blockade (Figure 3E). Furthermore, TRN inactivation increased the proportion of bimodal neurons and No-Go-responsive cells in MGB, while significantly reducing the fraction of stimulus-inhibited neurons (Figures 3F-3I, S7).

Together, these results show that auditory thalamus suppression by TRN is required by expert animals for both behavioral performance and to maintain single-cell coding specificity and stimulus-inhibited responses.

### TRN stabilizes population coding geometry in auditory thalamus

To determine how the single-cell changes observed upon TRN blockade could impair behavior, we analyzed population coding in MGB activity (Figure 4A). We first trained within-session population decoders to classify trial identity (Go/No-Go) from MGB activity (Figure 4B). Decoder accuracy was comparable across naïve, expert control and CNO sessions, indicating that stimulus separability in auditory thalamus is preserved across learning and after TRN blockade.

**Figure 4.**
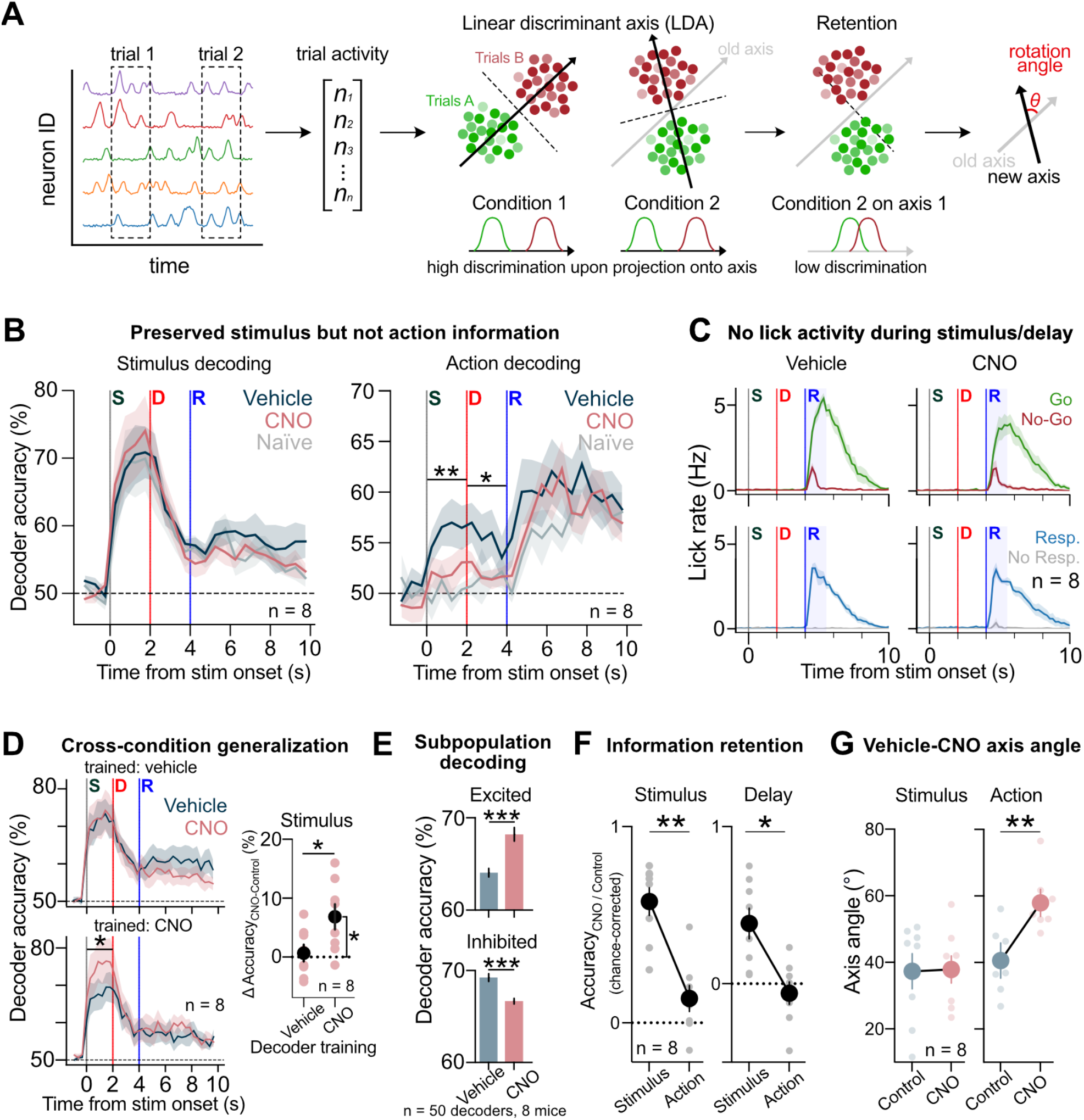
TRN blockade preserves stimulus information but alters population geometry. (**A**) Population geometry analysis. Trial activity was represented as N-dimensional population vectors, with each trial type (*e*.*g*., Go or No-Go) forming distinct point clouds. Linear Discriminant Axis (LDA, black solid arrow) identified axis separating trial types. Separability was quantified by projecting trials onto same-session axes, retention by projecting CNO activity onto the control-derived LDA axis and geometry changes by the angle between control and CNO axes. (**B**) Time-resolved balanced decoding of stimulus type (Go/No-Go; Sound/Visual) and action (lick/no-lick). Stimulus decoding was similar between control and CNO sessions, whereas action decoding was reduced by TRN blockade during stimulus and delay periods (Wilcoxon, stimulus p = 0.0078, delay p = 0.0234; n = 8), with similar response period accuracy. (**C**) Licks detected in video data, for Go/No-Go trials (top) and responded/non-responded trials (bottom). Tongue motion was virtually absent during stimulus and delay epochs and appeared only during the response period. Shaded area indicates the time window when mice had to lick for a reward. (**D**) Control-trained decoders generalized across conditions with above-chance accuracy, but trended lower in CNO late in the trial. CNO-trained decoders showed higher stimulus-period accuracy in CNO than control (Wilcoxon, p = 0.0156; n = 8). ΔCNO (Accuracy_CNO_ – Accuracy_Control_) was positive only for CNO-trained decoders (one sample Wilcoxon, p = 0.0156) and differed from control-trained decoders (Wilcoxon, p = 0.0078, n = 8). (**E**) Subpopulation decoding showed increased accuracy for stimulus-excited cells and decreased accuracy for stimulus-inhibited cells (50 decoders/mouse; Wilcoxon, both p < 0.001; threshold: 2xSD_baseline_). (**F**) Cross-condition retention, defined as chance-corrected CNO accuracy relative to control, was higher for stimulus-type than action coding during stimulus and delay periods (paired permutation: stimulus p = 0.0081; delay p = 0.0215; n = 8). (**G**) The trial-type (stimulus) discrimination axis rotated similarly across control days (vehicle and no injection) and across vehicle and CNO injection (analysis period: stimulus and delay, 0-4 s). However, the action axis (analysis period: stimulus and delay, 0-4 s, see also Fig. 4B) showed a greater rotation in the CNO session than across control sessions (paired t test, p = 0.0054, n = 8), indicating selective disruption of action-related population geometry under TRN blockade.

Because sensory thalamus can also encode behavioral choice^22^, we next asked whether MGB population activity predicted the animal’s upcoming action. Action decoding, defined as lick versus no-lick classification, was higher in expert control sessions than upon TRN silencing during both the stimulus (0–2 s) and delay (2–4 s) epochs (Figure 4B). Thus, in expert control conditions, upcoming actions are reflected in MGB activity before the response period, whereas this action-related signal is reduced by TRN blockade. This effect was not explained by overt licking during these periods (Figure 4C) and action decoding was not present in naïve animals (Figure 4B). Thus, these results indicate that TRN blockade does not eliminate stimulus information in MGB, but disrupts the population activity patterns that predict learned behavioral outcome.

We next tested whether TRN blockade altered the structure of the population code by assessing cross-condition generalization. Trial identity (Go/No-Go) decoders trained on expert control sessions generalized to CNO sessions (Figures 4D, S5A). In contrast, decoders trained on CNO sessions generalized less effectively to control sessions and performed significantly worse when tested on control data (Figure 4D). This asymmetric generalization indicates that TRN blockade induces a state-dependent reorganization of MGB population activity. Consistent with this interpretation, reduced cross-condition generalization was also observed during the response period (4–10s, Figure S5B).

To identify the cellular contributions to this reorganization, we next performed subpopulation decoding using stimulus-excited and stimulus-inhibited neurons separately. Compared to control conditions, TRN blockade increased decoding accuracy in stimulus-excited neurons but decreased it in stimulus-inhibited neurons (Figure 4E), revealing an imbalance between excitatory and inhibitory contributions to the population code. This population-level imbalance is consistent with the single-cell disinhibition observed under TRN blockade (Figure 3H).

We then directly examined whether this reconfiguration altered population coding geometry by quantifying cross-condition retention along the control-derived axis. For trial-type coding, CNO activity retained 62 ± 7% of the control above-chance decoding performance along this axis (Figure 4F), indicating that stimulus information remained present but was only partially aligned with the learned control axis. In contrast, action coding was more profoundly impaired by TRN blockade, retaining only 12 ± 6% of control performance along the same axis (Figure 4F). Consistent with this reduced retention, the discriminant action axis rotated between control and CNO conditions, beyond daily drift (57.95 ± 3°; Figure 4G). In addition, representational similarity analysis across control and TRN blockade conditions showed that trial-type representations became progressively less selective across trial epochs, particularly during the response period (Figures S6A, S6B). These findings are consistent with a degradation of the MGB population activity structure required for downstream behavioral readout.

Together, these results indicate that bottom-up sensory information remains decodable in auditory thalamus after TRN blockade, but that TRN-mediated inhibition is required to maintain the learned population coding geometry that links sensory representations to action. Loss of TRN-mediated inhibition therefore disrupts learned sensory behavior not by eliminating stimulus information, but by reconfiguring the MGB population code in a way that impairs reliable downstream readout.

### TRN tunes MGB toward reward-predicting cues

The preceding analyses showed that TRN blockade preserves stimulus information but disrupts the learned population geometry that links MGB activity to action. We next asked whether this TRN- and learning-dependent geometry reflects an acquired value bias in auditory thalamus. Comparing naïve and expert Go/No-Go sessions revealed bidirectional plasticity in MGB: responses to reward-predicting Go cues increased with learning, whereas responses to non-rewarded No-Go cues were suppressed (Figures 5A-5B, S8). Thus, experience reshaped thalamic cue representations according to learned stimulus value.

**Figure 5.**
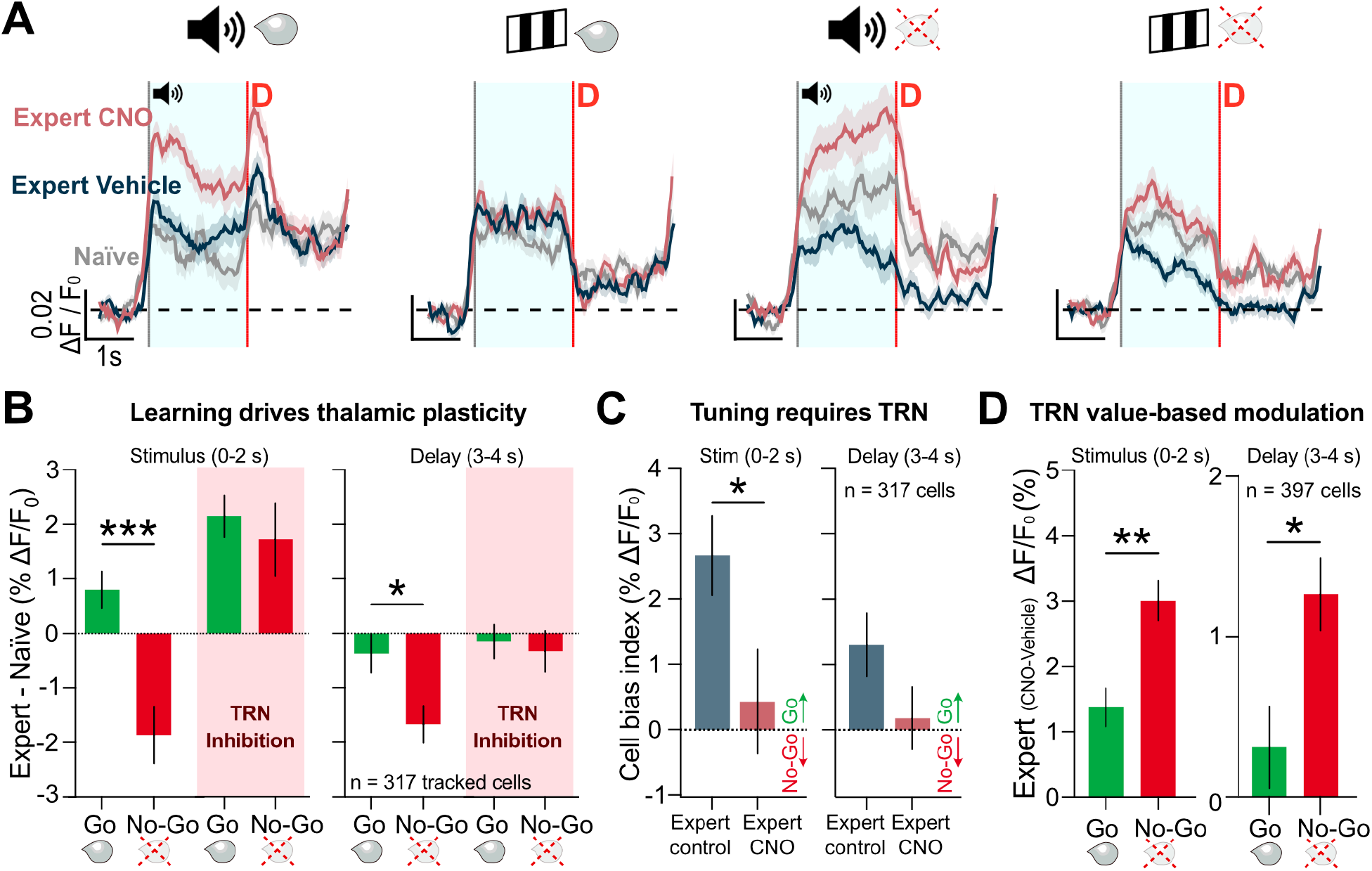
TRN enforces learning-dependent, value-based tuning in auditory thalamus. (**A**) Mean MGB population responses to Go-sound, Go-visual, No-Go-sound and No-Go-visual stimuli across naïve, expert control and TRN blockade (n = 317 tracked cells). (**B**) Learning-dependent plasticity, calculated as the change from naïve to expert for Go and No-Go stimuli (modalities combined) during stimulus and delay periods. Learning increased Go responses and suppressed No-Go responses (Mann-Whitney U: stimulus p < 0.001, delay p = 0.0181; n = 317 tracked cells, 9 mice), an effect abolished by TRN blockade. (**C**) Cell bias index (Go minus No-Go learning-dependent change) was reduced under TRN blockade (Mann-Whitney U, stimulus p = 0.0359; n = 317 cells). (**D**) TRN modulation strength (CNO – control) during expert performance. TRN blockade enhanced responses more strongly for No-Go than Go trials during stimulus (0-2 s) and delay (3-4 s) periods (Mann Whitney U; stimulus p = 0.0025, delay p = 0.0103; n = 397 cells, 9 mice), indicating stronger inhibition of non-rewarded cues under control conditions.

TRN blockade abolished this value-dependent regulation. In expert mice, suppression of No-Go responses was lost under TRN blockade (Figures 5B-5C, S8C-S8D, S8G-S8H). Moreover, delay-period suppression of No-Go activity, evident in expert controls, was likewise eliminated during TRN blockade (Figures 5B-C, S8G-S8H). To quantify this effect at the single-cell level, we computed a cell bias index defined as the learning-dependent change in Go responses minus the learning-dependent change in No-Go responses. TRN blockade lowered the bias index to near zero (Figure 5C), indicating a loss of differential tuning according to learned cue value. Direct comparison of TRN modulation across trial types further revealed stronger inhibitory control during No-Go than Go trials, both in stimulus and delay periods (Figure 5D). Thus, under expert conditions, TRN preferentially suppresses non-rewarded responses in MGB, enforcing a bias of sensory thalamic representations toward reward-predicting cues.

Together, these findings demonstrate that TRN dynamically supports learning-dependent value coding in auditory thalamus. By selectively suppressing non-rewarded signals while permitting or enhancing reward-predictive responses, TRN aligns thalamic cue representations with behavioral relevance and learned value (Figure 6).

**Figure 6.**
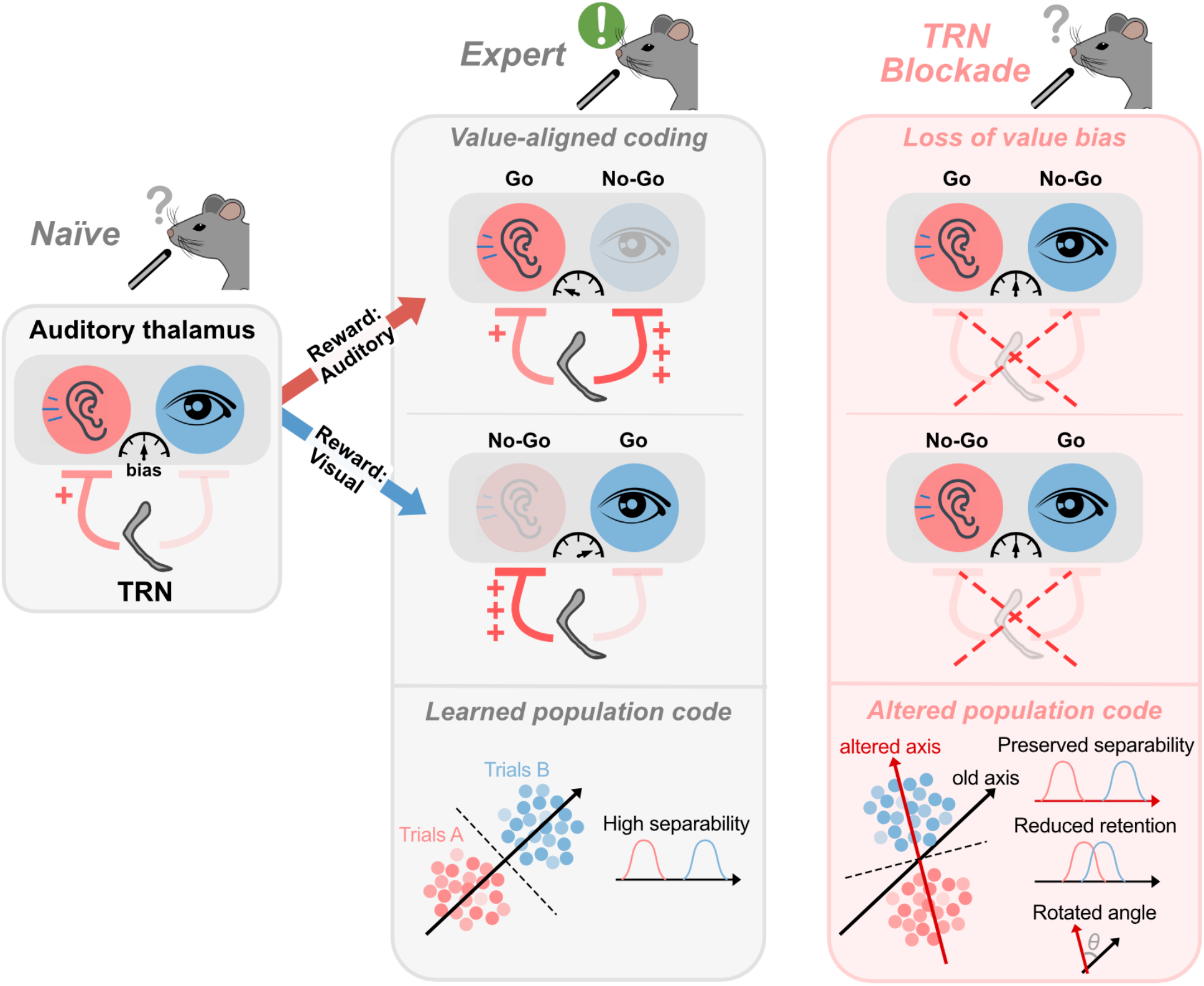
Model of TRN-dependent value alignment in auditory thalamus. In naïve animals, auditory thalamus exhibits a baseline sensory bias under ongoing inhibitory control from the thalamic reticular nucleus (TRN). Auditory cues are more strongly suppressed than visual stimuli. With learning, this baseline representation is reshaped into a value-aligned code. In expert animals, responses are biased towards the reward-predicting Go cue and suppressed for the non-rewarded No-Go cue, regardless of whether the rewarded cue is auditory or visual. This learning-dependent bias is mediated by TRN, which preferentially suppresses activity evoked by behaviorally irrelevant or non-rewarding stimuli, thereby aligning thalamic representations with learned stimulus value. TRN blockade abolishes this value-dependent tuning, leading to loss of bias and a more uniform representation of Go and No-Go stimuli. At the population level, expert control conditions are associated with a learned population code that is altered upon TRN blockade: despite preservation of stimulus separability, the retention of activity along the original control discriminant axis is reduced and the axis is rotated. Thus, TRN does not determine whether sensory information is present in auditory thalamus, but whether it is organized in a value-aligned manner that supports appropriate behavioral readout and action selection.

## Discussion

Adapting neuronal responses to sensory stimuli through experience is a core feature of the central nervous system. By filtering irrelevant signals, animals can conserve attentional and computational resources^16,23–25^, while strengthening responses to relevant cues facilitates their detection and the expression of learned behaviors^1,2,26^. Here we demonstrate that auditory thalamus flexibly adapts to behavioral demands through dynamic inhibition from the thalamic reticular nucleus (TRN). Blocking TRN-mediated inhibition altered sensory responses in auditory thalamus to both naïve and familiar cues, impaired task performance and revealed preferential TRN-dependent suppression of non-rewarded No-Go cues over reward-predicting Go cues. These findings identify inhibition from TRN as a circuit mechanism that aligns population activity with learned value to guide behavior.

Previous studies have implicated TRN in arousal^27^, sleep^28^ and attentional selection^16,25^. Our results extend this framework by identifying a learning-dependent role for TRN in shaping sensory thalamic coding according to acquired value. Rather than acting as a fixed gate for sensory throughput, TRN-mediated thalamic inhibition was dynamically shaped by learning history: it preferentially suppressed non-rewarded cues, enforced a bias toward reward-predicting stimuli and maintained action-related structure in MGB population activity. TRN thus emerges as a control center for learned relevance and value integration in thalamic computation.

Importantly, TRN blockade did not abolish stimulus information, as within-session stimulus decodability was preserved. This raises a central question: why does behavior fail while stimulus identity remains decodable? Preserved decodability by an *in silico* machine learning-based classifier does not necessarily mean that information remains equally accessible to the animal’s downstream readout circuits. Indeed, cross-condition analyses revealed reduced preservation of population structure, rotation of coding axes and particularly strong disruption of action-related coding. These findings suggest that TRN stabilizes a learning-dependent population code required for reliable behavioral readout. In this view, auditory thalamus continues to represent sensory information under TRN blockade, but this information is no longer embedded in the value- and action-aligned geometry established during learning^29-31^. Without TRN, MGB activity shifts into a different coding regime: new excitatory responses emerge, stimulus-inhibited populations are reduced, bimodal responses increase and the learned value-based bias is lost. This more diffusely excited sensory representation may remain informative to an external decoder, yet be poorly matched to downstream circuits trained by experience to interpret the TRN-gated code. Appropriate behavior therefore breaks not because sensory information is absent, but because its alignment with action-related readout is perturbed. Candidate downstream circuits mediating this readout include sensory cortex^6^, fronto-thalamocortical loops^32^ and striatal/amygdala^1,33^ circuits. Future work will be required to determine how these sub-networks interpret value-aligned thalamic representations to guide behavior.

Moreover, upon TRN blockade, MGB activity was amplified not only during stimulus presentation but also during the delay and response periods (Figures S8-S9), indicating that TRN shapes thalamic population dynamics across the entire trial timeline. Thus, inhibitory control of auditory thalamus by TRN contributes not only to sensory encoding but also to decision-related processes such as stimulus memory, action selection or reward expectation^2,34^. Future work will determine how rapidly TRN adapts upon shifts of cue-outcome contingencies or contextual rule changes, revealing the flexibility of higher-order inhibitory control over thalamic processing.

We also uncovered modality asymmetries in TRN-mediated control of MGB. Auditory stimuli were consistently modulated, regardless of predictive value, whereas visual stimuli were suppressed primarily when they signaled the absence of reward. This suggests the existence of specialized TRN subnetworks targeting distinct sensory thalamic nuclei. Because our recordings targeted auditory thalamus, auditory stimuli may be particularly sensitive to TRN-mediated control, including when they serve as Go cues. Future experiments mapping TRN subnetworks across sensory domains will clarify whether such asymmetries generalize across modalities^10,35^.

TRN also modulated responses to novel stimuli in naïve animals, indicating a role in general perceptual regulation. Moreover, TRN suppression reduced the proportion of stimulus-inhibited cells in MGB. This neuron subset may sharpen selectivity, support top-down control, or encode information complementary to excitatory ensembles^36–42^. Their loss may impair perceptual precision without causing a gross loss of stimulus information. Thus, TRN may contribute not only to the filtering of familiar, task-relevant signals, but also to the fidelity of perceptual representations.

Our results further raise the question of how TRN plasticity is itself instructed. Higher-order regions – including prefrontal cortex, auditory cortex, basal ganglia and amygdala – provide convergent inputs to TRN and may regulate its activity according to task rules^43^, reward associations^44–45^ and internal states^46^. Through this convergence, TRN may align sensory thalamic coding with task rules, motivational context and internal state.

Finally, our results have translational implications for human diseases. Disruptions of TRN-thalamus circuits are implicated in schizophrenia, autism spectrum disorder and ADHD – conditions characterized by aberrant sensory gating and impaired salience attribution^3,47–50^. Sensory overload and inflexible behavioral responses may arise from failures in such inhibitory regulation^51^. By revealing how TRN dynamically gates thalamic activity according to stimulus history and learned value, our work provides a mechanistic framework for understanding these deficits. More broadly, TRN emerges as a dynamic regulator of thalamocortical communication, ensuring that sensory signals are represented in accordance with learned experience, value and behavioral demands.

## Methods

### Animals

All experiments were performed in accordance with the institutional guidelines of the University of Bonn and DZNE and approved by the Landesamt für Natur, Umwelt und Verbraucherschutz of North Rhine-Westphalia, Germany. Adult male and female PV-Cre mice (B6.129P2-Pvalb^tm1(cre)Arbr^/J, The Jackson Laboratory) were used. Mice were housed under a 12 h light/dark cycle at controlled temperature (22°C) and humidity (55%), with full access to food and water before behavioral experiments. During training, food was restricted to maintain body weight at 85-90% of baseline *ad libitum* weight, and animals’ well-being was monitored daily.

### Surgical procedures

Mice were anesthetized with isoflurane (induction 5%, maintenance 1.5-2%) delivered in oxygen-enriched air (95%, 1-3 L/min, Oxymat III, Weinmann). Tail and foot reflexes were tested prior to surgery. Animals were positioned in a stereotaxic frame (Model 1900, Kopf Instruments), and body temperature was maintained at 36.5 – 37 °C with a heating pad (Rodent warmer, 53800 M, Stoelting). Eyes were protected with eye cream, and a local anesthetic mixture of lidocaine (10 mg/kg) and ropivacaine (3 mg/kg) was injected subcutaneously into the scalp.

Craniotomies were made with a stereotaxic drill (Model 1911, Kopf) and 0.3 mm microdrill bits (105-0118.225, Kyocera): one above the medial geniculate body (MGB; AP: −3.28, ML: −1.9, DV: 3.2 mm) and two above the thalamic reticular nucleus (TRN; AP: −1.7, ML: ±2.2, DV: 3.2 – 3.6 mm). Viral vectors were injected through pulled glass pipettes (2-000-001, Drummond Scientific) using a pressure ejection system (Picospritzer, Parker). Vectors ssAAV-1/2-hSyn1-dlox-hM4D(Gi)-mCherry(rev)dlox-WPRE-GHp(A) and ssAAV-2/1-hSyn1-dlox-mCherry(rev)-dlox-WPRE-hGHp(A) were a gift from Bryan Roth^54^ and were injected in the TRN of experimental and control groups, respectively. ssAAV-5/2-mCaMKIIα-jGCaMP8m-WPRE-bGHp(A) was a gift from the GENIE Project^55,56^ and was injected into MGB. Viral vectors were obtained from the Viral Vector Facility Zurich of the Neuroscience Center Zurich (ZNZ) and diluted 1 – 2 x in sterile PBS, and 300-400 nL were injected. TRN injections were delivered in steps along the dorsoventral axis (DV: 3.6, 3.4, 3.2 mm). Drill and pipette positioning was performed with a micropositioner (Model 2650, Kopf).

One week later, mice underwent a second surgery for implantation of a gradient refractive index (GRIN) lens (0.7 mm diameter, 7.3 mm length, Inscopix) above MGB, together with a custom titanium head bar. Both were fixed to the skull with curable glue (Loctite 4305, Henkel). To prepare for lens placement, a 0.78 mm craniotomy was made (226-0307.400, Kyocera), and a track was created with a sterile 0.7 mm needle. The skull was sealed with Scotchbond (3M), Vetbond (3M) and dental cement (Paladur). Animals received systemic analgesia via carprofen (Rimadyl, Zoetis, 0.067 mg/mL in drinking water) from 24 h before to 72 h after surgery.

### Two-photon *in vivo* calcium imaging and behavior

A custom-built two-photon microscope (INSS) with a resonant scanning system and a pulsed Ti:sapphire laser (λ = 940 nm, Chameleon Vision S, Coherent) was used for all imaging experiments. Positioning and focus on the GRIN lens were achieved using a motorized three-axis system (Zaber motors), which allowed movement of the microscope objective (Z direction) and the behavioral apparatus breadboard (X–Y directions). The microscope was controlled via ScanImage software (Vidrio Technologies, MBF Bioscience). Fluorescent photons were collected with a 16x, 0.80 NA objective lens (Nikon), split by a dichroic mirror (T565lpxr, long pass, Chroma) and passed through barrier filters (green: ET510/80 m, red: ER630/75 m) before detection with PMTs (PMT2101, Thorlabs). Imaging was performed at 512 x 512 pixels and 30 Hz frame rate.

During imaging and behavior, mice were head-restrained with a custom holding system and placed on a running wheel connected to a rotary encoder (E6A2–CS3E, Omron). A custom 3D-printed tunnel was placed above the animal to cover the body and minimize stress. Auditory stimuli (pure tones) were presented via a speaker (ES1, Tucker Davis Technologies) positioned 10 cm away at the contralateral upper-right side of the mouse. Visual stimuli (moving gratings on a continuous gray background) were displayed on a 7-inch screen (Adafruit 1667) positioned 10 cm from the contralateral side of the face.

For behavioral testing, a custom-made lick spout mounted on a motorized linear stage (Zaber) approached the snout during the “response” window of each trial, allowing the mouse to report. Correct responses (“hits”) triggered reward delivery through a custom lick-detection system coupled to a peristaltic micropump (mp6, Bartels Microtechnik), which dispensed an 8-10 μL droplet of vanilla-flavored soy milk.

All experiments were controlled by a custom MATLAB (MathWorks) software using the Psychophysics Toolbox (http://psychtoolbox.org), in combination with a multifunction data acquisition device (USB-6008, National Instruments) and a real-time signal processor (RZ6, Tucker Davis Technologies). Behavioral events were synchronized with imaging via TTL signals recorded at 50 Hz.

### Behavioral training and analysis

Mice were habituated to head fixation in the recording setup for at least 3 days. In addition, each behavioral session began with a 1-min habituation period.

For naïve passive-exposure experiments, auditory (4, 8, 12, 16, 20 kHz pure tones, 75 dB), visual (upward, rightward, downward and leftward drifting gratings; 100% contrast, 2 Hz, 0.05 cycles/degree) and multisensory (all possible unisensory combinations) stimuli were presented in pseudo-random order. Each trial type was repeated eight times (240 total stimuli per session). Stimuli lasted 2 s, with an intertrial interval (ITI) of 6–9 s. Passive-exposure sessions were conducted 45–60 min after intraperitoneal injection of either CNO (clozapine N-oxide dihydrochloride, Tocris; 3 mg/kg in 0.3 mL Mill-Q water) or vehicle (carrier solution).

For the Go/No-Go sensory detection task, mice first underwent 1–2 sessions of lick training without stimuli to habituate them to the lick-reward system. Animals were then trained on a Go/No-Go task with two stimulus types: 12 kHz tones and rightward-drifting gratings. Sound-trained mice were rewarded for reporting tone trials and required to withhold licking to gratings; visual-trained mice followed the inverse rules. Both versions included catch trials (no stimulus), which always had to be ignored. Task composition was 30% Go, 35% No-Go and 35% catch trials (140 total trials per session). Each trial consisted of a 6–13 s ITI, followed by a 2 s stimulus presentation, a 2 s delay, and a 1.5 s response window, during which the lick spout was positioned ~ 1 cm from the snout and retracted at the end.

Mouse licking was captured in the response periods by the lick spout sensor. Licks outside the response period were detected via DeepLabCut 3.0^52^, but were extremely rare (Figure 4C). Behavioral performance was quantified using hit rate (correct Go trials / total Go trials) and false alarm rate (incorrect responses on No-Go and catch trials / total No-Go and catch trials). Detection sensitivity, independent of response bias^53^, was assessed using d’ (sensitivity index), defined as the difference between the z-transformed hit rate and false alarm rates: *d’ = z(hit rate) – z(false alarm rate)*. Higher d’ values indicate better discrimination between Go and No-Go/catch trials. Mice were considered expert after reaching d’ > 1–1.5 in two consecutive sessions. Expert animals subsequently underwent vehicle and CNO injection sessions. Seven of nine mice received vehicle first, while two of nine received CNO first. Results were consistent across injection orders, and CNO-induced performance impairments were acute, as animals returned to baseline performance 24 h post-injection.

### Two-photon image processing and analysis

Two-photon images were processed using Suite2p^57^. Motion correction was applied, and regions of interest (ROIs) were automatically detected. ROIs were then manually curated: neurons were selected, non-neuronal ROIs were discarded and missing cells were added with the manual ROI drawing tool in Suite2p. Each neuronal trace, together with its neuropil signal, was exported for downstream analysis in Python.

Neurons were manually tracked across imaging sessions for each mouse using maximum-intensity projection images. To facilitate longitudinal tracking, we used a custom head bar system that minimized angular rotation of the imaging plane, allowing repeated recordings from the same plane. Between 30–90 neurons were recorded per mouse.

For each neuronal trace, a correction coefficient (0.7) was multiplied with the neuropil signal, and the result was subtracted from the raw ROI signal. The resulting trace was considered the neuronal Ca^2+^ signal. Data were downsampled from 30 Hz to 5 Hz, aligned to trial onsets, and ΔF/F_0_ was calculated using the 1 s pre-stimulus period as baseline.

To generate population heatmaps, the average response for each cell and trial type was computed, and cells were sorted by response magnitude during the stimulus period (0–2 s). Analyses of specific subperiods are indicated in the figures and include: stimulus period (0–2s from stimulus onset), delay period (3–4s, excluding stimulus offset responses) and combined stimulus and delay period (0–4s).

For response subtype analysis, cells were classified as excited if their stimulus-period activity exceeded 2 x baseline standard deviation (SD), and as inhibited if activity fell below – 2 x baseline SD. For reward response analysis, traces were aligned to reward delivery times.

Learning-dependent response change was calculated for each tracked neuron as the difference between expert vehicle and naïve session responses within the defined analysis window (stimulus period: 0-2 s; delay period: 3-4 s). For Figure 5 analyses, responses were averaged across auditory and visual stimuli according to trial type. A cell bias index was computed for each neuron as the following: Bias Index = ΔR_Go_ – ΔR_No-Go_; where ΔR represents the learning-dependent change (expert vehicle-naïve). Positive values indicate stronger learning-related enhancement for Go relative to No-Go cues. Bias indices were compared between vehicle and CNO conditions to assess TRN-dependent modulation of value-based tuning.

In the multisensory integration analysis, a linearity index was calculated as: Linearity index = R_Multi_ – (R_Uni1_ + R_Uni2_); where R_Multi_ is the response to the multisensory stimulus and R_Uni1_ and R_Uni2_ are the corresponding unisensory responses. Negative values indicate sublinear integration.

### Population and decoder analysis

Decoding analysis was performed using custom Python implementations using NumPy. SciPy and Decodanda^58^.

For same-session decoding, neural population vectors were computed by averaging activity within defined time windows, and linear decoders were constructed using a shrinkage-regularized linear discriminant analysis (LDA) approach. To decode behavioral outcome (lick vs no-lick), trials were labeled based on the animal’s response and pooled across stimulus conditions. A binary LDA classifier was trained to discriminate lick versus no-lick trials using cross-validation with repeated split-half resampling. Decoder performance was quantified using balanced accuracy on held-out trials to account for class imbalance.

To assess cross-condition generalization, decoders trained in one condition (e.g., vehicle) were applied to data from another condition (e.g., CNO), and generated a single decoding value for the entire period (*e*.*g*., stimulus or delay). A retention index was computed as the ratio of decoding performance in the test condition relative to the training condition, after chance-correction: Retention = (Accuracy_test_ - 0.5) / (Accuracy_train_ - 0.5).

For geometric analyses, population coding axes were defined as the normalized LDA weight vectors separating task variables (stimulus: Go vs No-Go, action: lick vs no-lick). The angle between axes across conditions (e.g., vehicle vs CNO) or variables (stimulus vs action) was computed using the arccosine of the dot product between normalized axes, providing a measure of representational alignment or rotation in population space. To account for finite-trial sampling variability in axis estimation, cross-condition angles were corrected by subtracting the mean within-session axis rotation estimated from the two sessions being compared. Within-session axis rotation was estimated by repeatedly fitting LDA axes to random 75%-trial subsamples and measuring the angle between each subsampled axis and the corresponding full-session axis.

For cross-session, time-resolved decoding, as well as for subpopulation decoding, Decodanda was used. The decoder was trained to classify trial identity (sound vs. visual) using 70% of the data from both sound- and visual-trained mice in their last expert (no-injection) session. Performance was evaluated on the remaining 30% hold-out trials, as well as on vehicle and CNO session data to assess cross-session generalization.

For time-resolved, intra-trial decoding, classification accuracy was computed in 100 ms bins spanning from 1 s before to 10 s after stimulus onset. Accuracy values were first computed per mouse and then used for statistical comparisons.

To quantify state-dependent changes in decoding performance, a CNO modulation index (ΔCNO) was computed for each mouse and time bin as: ΔCNO = Accuracy_CNO_ – Accuracy_Vehicle_ (Figure 4), or ΔCNO = Accuracy_CNO_ – (Accuracy_Vehicle_ + Accuracy_No-injection_) / 2 (Figure S5). For comparisons between baseline-trained and CNO-trained decoders, ΔCNO values were computed separately for each decoder type. Statistical analyses were performed on per-mouse ΔCNO values.

For subset analyses of stimulus-excited and stimulus-inhibited neurons, decoding was performed using a 1 s window corresponding to the second half of the stimulus period (1-2 s after stimulus onset). Cells were classified as excited or inhibited using the same 2 x baseline SD criterion described above. To avoid trial- or cell-specific biases, 50 independent decoding models were trained per mouse with randomized train-test splits. For each mouse, the number of neuron traces was balanced across conditions to ensure equal numbers of excited and inhibited cells were included.

Population-level representations were further analyzed using a representational similarity approach based on population vector correlations. For each session and epoch, neural activity was averaged across trials to obtain condition-specific population vectors (Go, No-Go and catch). Pearson correlation coefficients were then computed between population vectors across conditions (e.g., vehicle vs CNO), generating cross-condition correlation matrices.

To quantify the preservation of stimulus coding, correlations between matching trial types (e.g., Go-Go) were compared to correlations between mismatched trial types (e.g., Go-No-Go). A selectivity preservation index was defined as the difference between same-type and different-type correlations (same minus different), providing a measure of how well task-relevant population structure was maintained across conditions.

### Histology

At the end of experiments, mice were transcardially perfused with PBS followed by 40 mL, 4% paraformaldehyde (PFA) in PBS. Brains were extracted, post-fixed overnight at 4°C in 4 % PFA, and transferred to PBS. Coronal slices (150 μm) were cut with a vibratome (Leica VT1000 S) and immunostained for calretinin (MGB marker^1^) and parvalbumin (TRN marker).

Immunohistochemistry included: PBS washes, 2 h incubation at room temperature in blocking solution (10% normal horse serum, Histoprime LIN-ENH9010-10, Biozol; 0.5% Triton X-100, 11488696, Alfa Aesar), overnight incubation at 4°C with primary antibodies (goat anti-calretinin, 1:1000, CG1, Swant; guinea pig anti-parvalbumin, 1:500, 195308, Synaptic Systems) in carrier solution (1% horse serum, 0.5% Triton in PBS), washes in 0.5% Triton PBS, and 2 h incubation with secondary antibodies (donkey anti-goat Alexa Fluor 647, 1:1000, A-21447, Invitrogen; goat anti-guinea pig IgG Alexa Fluor 488, 1:1000, A11073, Invitrogen; DyLight 405-conjugated AffiniPure goat anti-guinea pig IgG, 1:250, 106-475-003 Jackson Immuno Research). Slices were washed, mounted on glass slides (Superfrost, BB02400600S113MNZ0, Epredia) with coverslips (X100 coverslips 24 x 60 mm, 0.13-0.16 mm, Epredia). Images were acquired with an LSM700 confocal microscope (Zeiss) and a fluorescence Axio Zoom v16 microscope (Zeiss).

### Statistics

Statistical analysis was performed in Prism 10 (GraphPad) and Python. The alpha level was set at 0.05 and multiple-comparisons corrections were applied when appropriate. Statistical tests and p values are reported in the text. Kolmogorov-Smirnov was used to assess normality of the data. Data are shown as mean *±* SEM unless otherwise stated. Statistical significance is indicated as p < 0.05 (*), p < 0.01 (**), and p < 0.001 (***).

## Acknowledgements

We thank Julia Esser for assistance with mouse breeding and histology, and Juliane Schiweck, Ziyan Huang and Sherman Richard Chau for helpful comments on the manuscript. This work was supported by the European Research Council starting grant 803870 (J.G.); the Deutsche Forschungsgemeinschaft (DFG, German Research Foundation) through SFB1089 (J.G.), SPP2411 (J.G.), Nahostkooperation 559314332 (J.G.), and Walter Benjamin grant 528405672 (R.P.M.); the iBehave Network (J.G.); Helmholtz AI; and the Deutsches Zentrum für Neurodegenerative Erkrankungen (DZNE).

## Contributions

R.P.M. and J.G. conceptualized the project and designed the experiments and methodology. R.P.M. acquired the data. R.P.M. and M.K. performed formal analyses. R.P.M., M.K. and A.B. designed analysis software and performed data curation. J.G. supervised the study. R.P.M. and J.G. wrote the original draft of the manuscript with input from all authors.

## Supplemental Information

**Suppl Figure 1.**
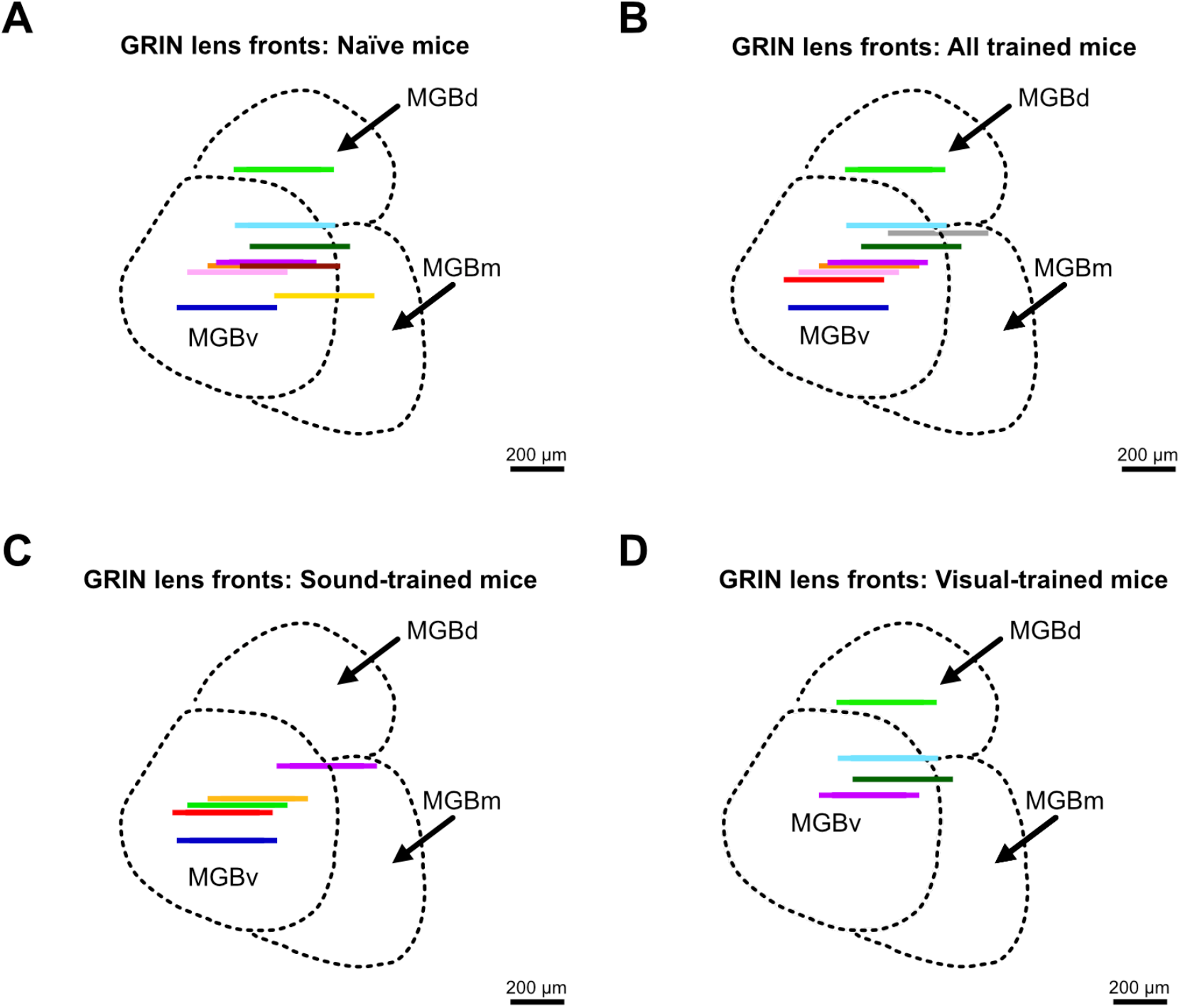
GRIN lens implantation sites in auditory thalamus (MGB). (**A**) GRIN lens front locations for mice used in the naïve passive CNO experiment (n = 9). (**B**) GRIN lens front locations for mice performing the Go / No-Go task and tested with CNO at the expert stage (n = 9; sound-trained and visual-trained combined). (**C**) GRIN lens front locations for sound-trained mice (sound = Go; n = 5). (**D**) GRIN lens front locations for visual-trained mice (visual = Go; n = 4).

**Suppl Figure 2.**
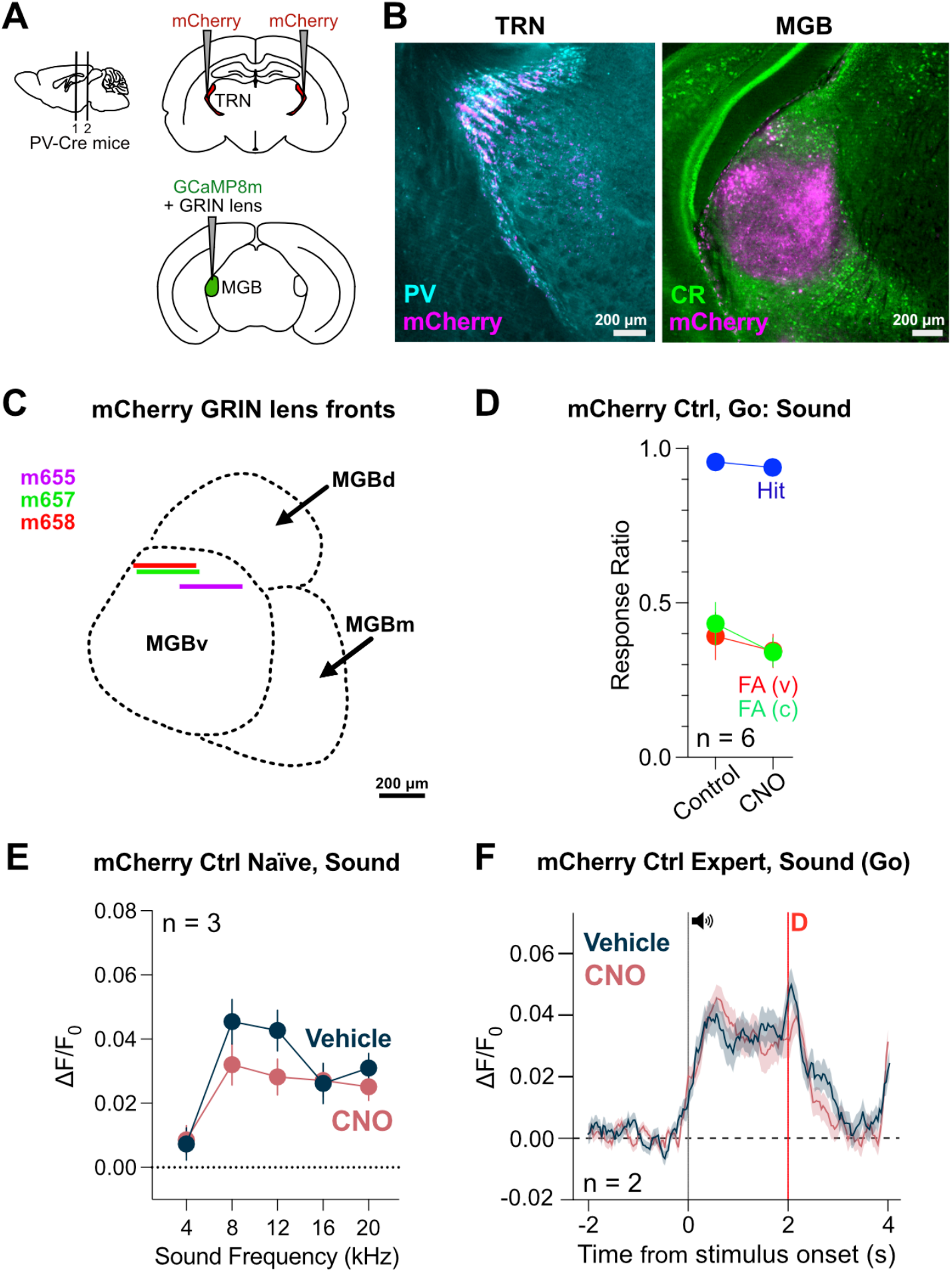
mCherry control mice show no behavioral or thalamic effects of CNO injection. (**A**) Control mice were bilaterally injected with mCherry (DREADD-negative control) in TRN and GCaMP8m in MGB. A GRIN lens was implanted above MGB. (**B**) Histological sections confirm mCherry expression in TRN and axonal projections to MGB. (**C**) GRIN lens front locations for mCherry-injected mice (n = 3). (**D**) In the Go/No-Go task (expert stage), hit and false-alarm rates were unchanged following CNO injection compared to vehicle. (**E**) In naïve passive sessions, CNO injection did not alter MGB auditory responses in mCherry control mice (n = 3). (**F**) In expert sound-trained mCherry control mice, CNO injection did not change mean MGB responses to the sound stimulus. Data show mean ± SEM. Statistical significance: *p<0.05, **p<0.01, ***p<0.001.

**Suppl Figure 3.**
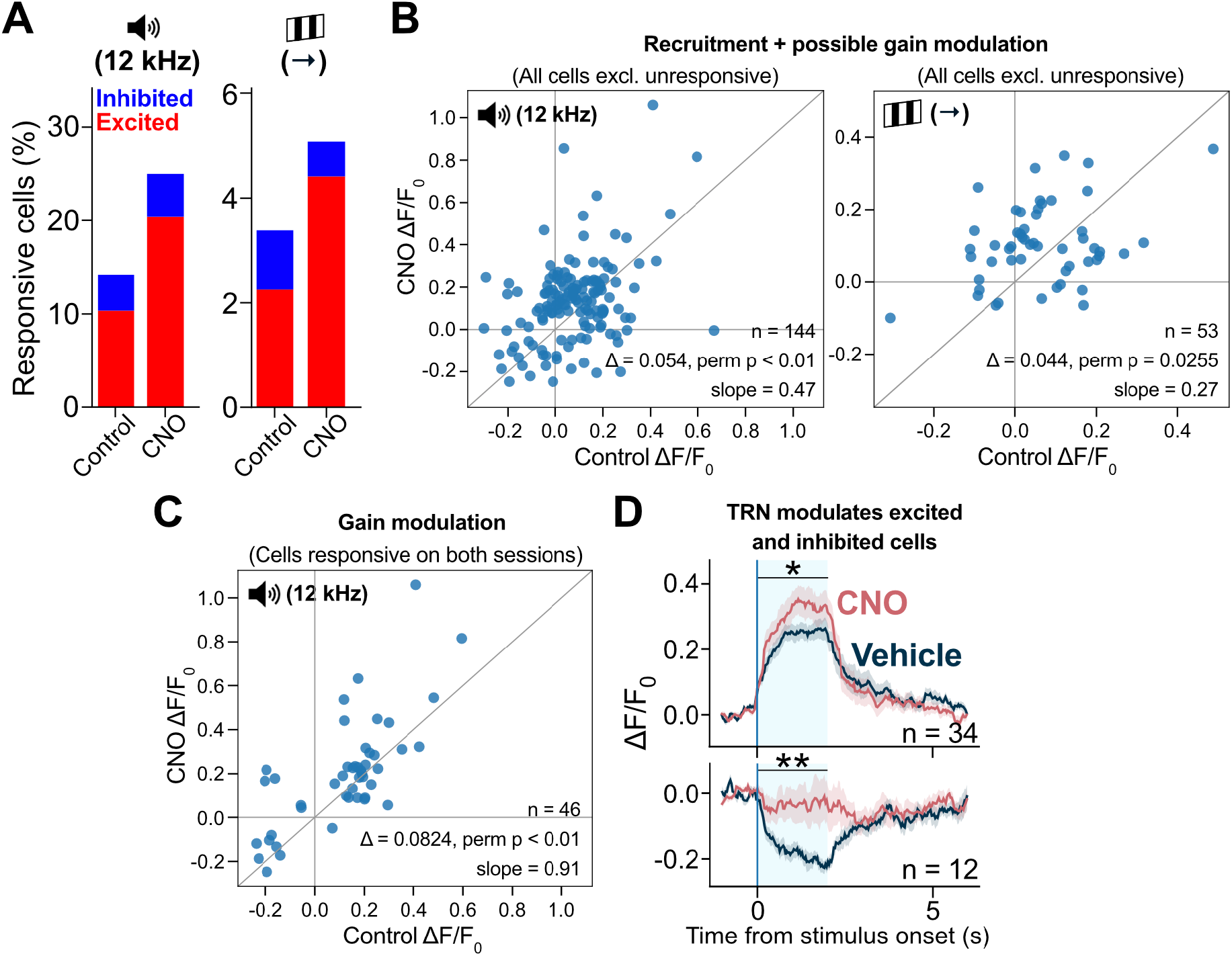
TRN blockade increases response recruitment and gain during naïve sensory processing. (**A**) Proportion of MGB cells responsive to sound (12 kHz) and visual (**→** grating) stimuli during passive exposure under vehicle and CNO conditions (n = 444 cells from 9 mice). (**B**) Scatter plots comparing individual cell responses to sound (left) and visual (right) stimuli in vehicle versus CNO sessions. Cells unresponsive in both sessions were excluded. Responses were significantly larger under CNO for sound (paired permutation test, p = 0.00105, n = 144 tracked cells) and visual stimuli (paired permutation test, p = 0.0255, n = 53 tracked cells), consistent with increased recruitment and/or response amplification. (**C**) Among neurons responsive to sound in both sessions, response amplitudes were greater under CNO than vehicle (paired permutation test, p = 0.00155, n = 46 tracked cells), indicating a gain modulation of already responsive cells. (**D**) PSTHs of sound-responsive neurons classified as excited or inhibited show enhanced excitation and reduced inhibition under CNO (permutation tests, excited cells p = 0.0391; inhibited cells p = 0.0051). Data show mean ± SEM. Statistical significance: *p<0.05, **p<0.01, ***p<0.001.

**Suppl Figure 4.**
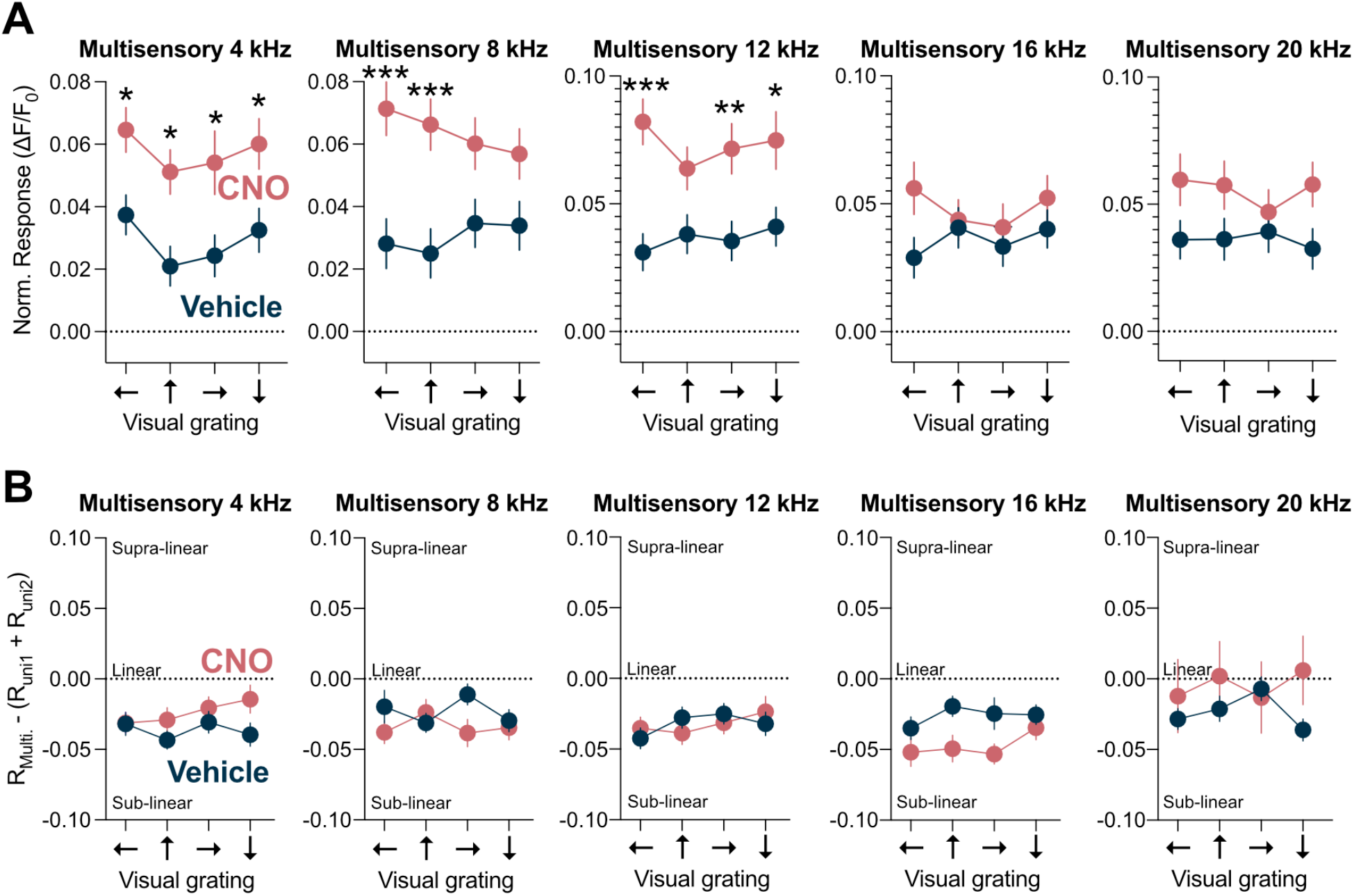
Multisensory integration remains sublinear under TRN blockade. (**A**) Mean MGB responses to multisensory combinations of auditory (4, 8, 12, 16, 20 kHz tones) and visual (leftward, upward, rightward, downward gratings) stimuli during passive exposure in naïve mice. Several stimulus combinations showed enhanced response amplitudes under TRN blockade (two-way ANOVA with Bonferroni correction; 4 kHz ⟵ p = 0.0393, 4 kHz ↑ p = 0.0167, 4 kHz **→** p = 0.0187, 4 kHz ↓ p = 0.0348; 8 kHz ⟵ p < 0.001, 8 kHz ↑ p < 0.001; 12 kHz ⟵ p < 0.001, 12 kHz **→** p = 0.0059, 12 kHz ↓ p = 0.0122; n = 414 cells from 9 mice). (**B**) Linearity index analysis revealed no significant change in multisensory integration under TRN blockade (n = 414 cells from 9 mice). In both control and CNO conditions, multisensory responses remained sublinear, indicating that the observed amplitude increases primarily reflect enhanced auditory drive rather than altered integration rules. Data show mean ± SEM. Statistical significance: *p<0.05, **p<0.01, ***p<0.001.

**Suppl Figure 5.**
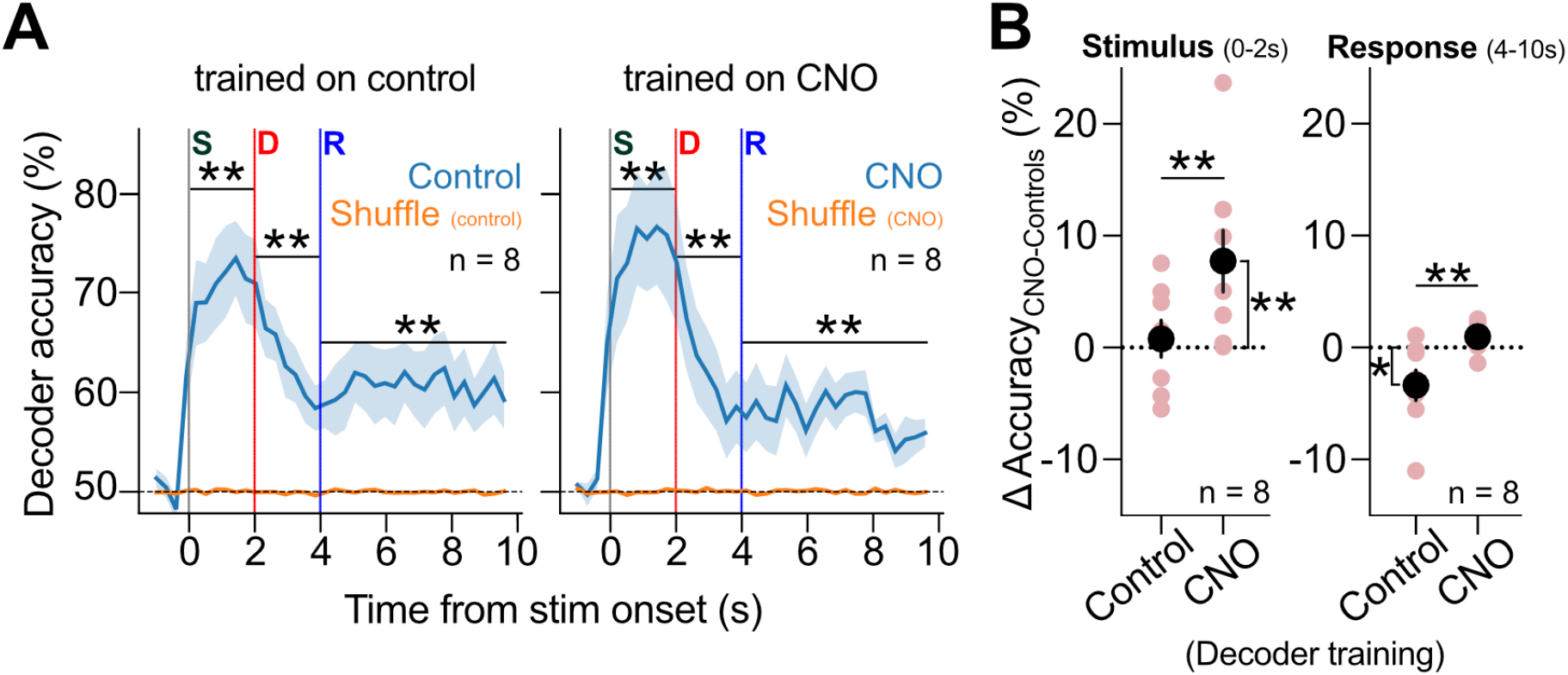
Supplementary Go/No-Go trial decoding data. (**A**) Decoder accuracy dropped significantly when shuffling trial identity (Wilcoxon, p = 0.0078 for stimulus 0-2 s, delay 2-4 s and response 4-10 s, for both control-trained and CNO-trained decoders, n = 8 mice). (**B**) For this analysis, ΔCNO was calculated taking into account two control sessions: vehicle injection control and no injection control: ΔCNO = Accuracy_CNO_ – (Accuracy_Control-no-injection_ + Accuracy_Control-vehicle_) / 2. Similar to Figure 3, ΔCNO (Accuracy_CNO_ – Accuracy_Controls_) during the stimulus (0-2 s) period was significantly greater than zero only for CNO-trained decoders (one sample Wilcoxon, p = 0.0078) and differed from control-trained decoders (Wilcoxon matched pairs, p = 0.0078, n = 8; left). During the response (4-10 s) period, ΔCNO was significantly lower than zero only for control-trained decoders (one sample Wilcoxon, p = 0.0391) and differed from CNO-trained decoders (Wilcoxon matched pairs, p = 0.0078, n = 8; right). Data show mean ± SEM. Statistical significance: *p<0.05, **p<0.01, ***p<0.001.

**Suppl Figure 6.**
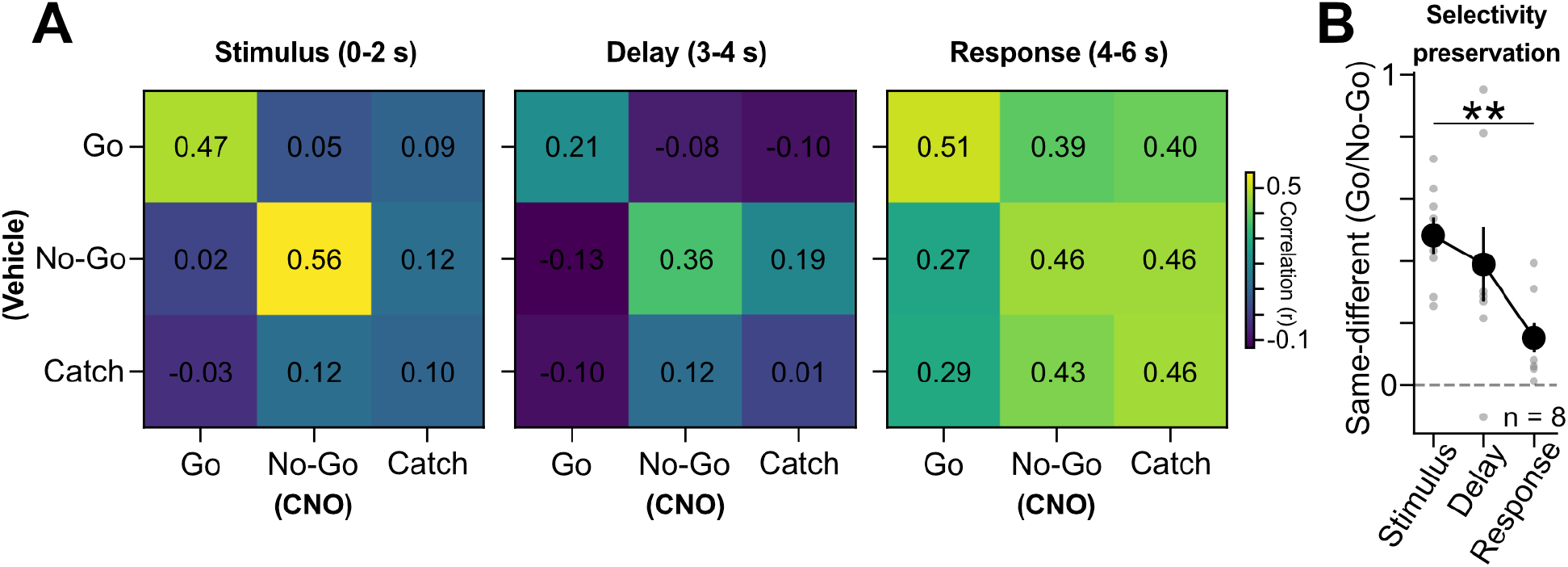
TRN blockade progressively reduces cross-condition selectivity of MGB representations. (**A**) Population vector correlation matrices between control and CNO sessions across trial types (Go, No-Go, catch) for stimulus (0–2 s), delay (3–4 s) and response (4–6 s) epochs. Matrices show mean pairwise correlations of population activity vectors, organized by trial type and condition. (**B**) Cross-condition coding selectivity, defined as the difference between correlations of same trial types and different trial types (same – different). Selectivity declines across epochs and is reduced in the response compared to the stimulus period (paired permutation test, p = 0.008; n = 8), indicating reduced preservation across conditions late in the trial. Data show mean ± SEM. Statistical significance: *p<0.05, **p<0.01, ***p<0.001.

**Suppl Figure 7.**
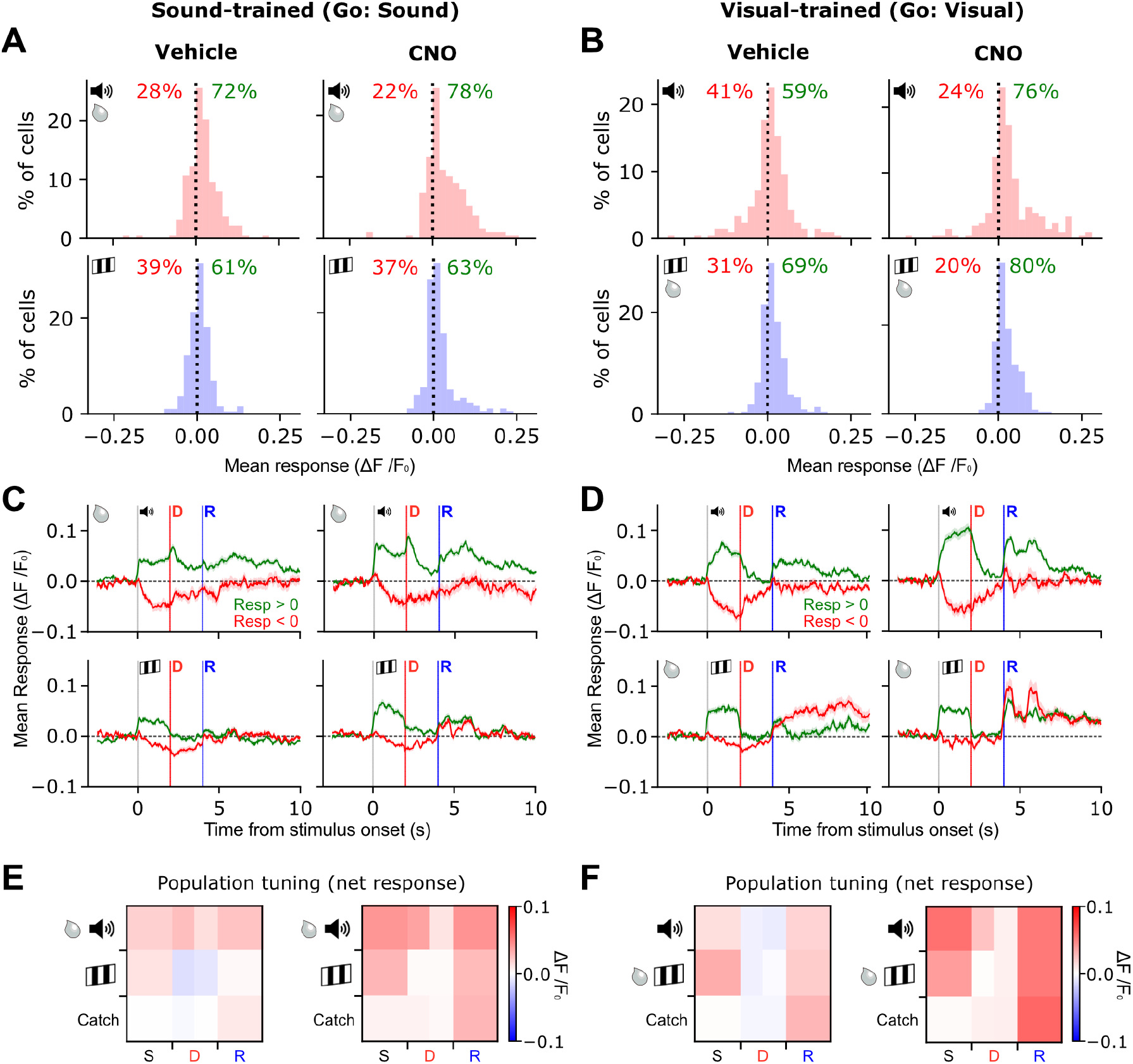
TRN blockade shifts excitatory-inhibitory balance and population dynamics in auditory thalamus. (**A**) Histograms of MGB cell responses to sound and visual stimuli in sound-trained expert mice (Go: sound), under vehicle (left) and CNO (right) conditions. (**B**) Same analysis as in (**A**) for visual-trained expert mice (Go: visual). (**C**) Mean population response PSTHs of stimulus-excited (green) and stimulus-inhibited (red) cells during Go sound trials (top) and No-Go visual trials (bottom) in sound-trained mice, under vehicle (left) and CNO (right). Stimulus (S), delay (D) and response/action (R) periods are indicated. (**D**) Same as in (**C**), but for visual-trained expert mice. (**E**) Mean MGB activity matrices for sound-trained mice (Go: sound) under vehicle (left) and CNO (right). Trial epochs are indicated as stimulus (S), delay (D) and response/action (R). The delay period is subdivided to separate stimulus offset responses. (**F**) Mean MGB activity matrices for visual-trained mice (Go: visual) under vehicle (left) and CNO (right). (n = 188 sound-trained cells, n = 209 visual-trained cells). Data show mean ± SEM.

**Suppl Figure 8.**
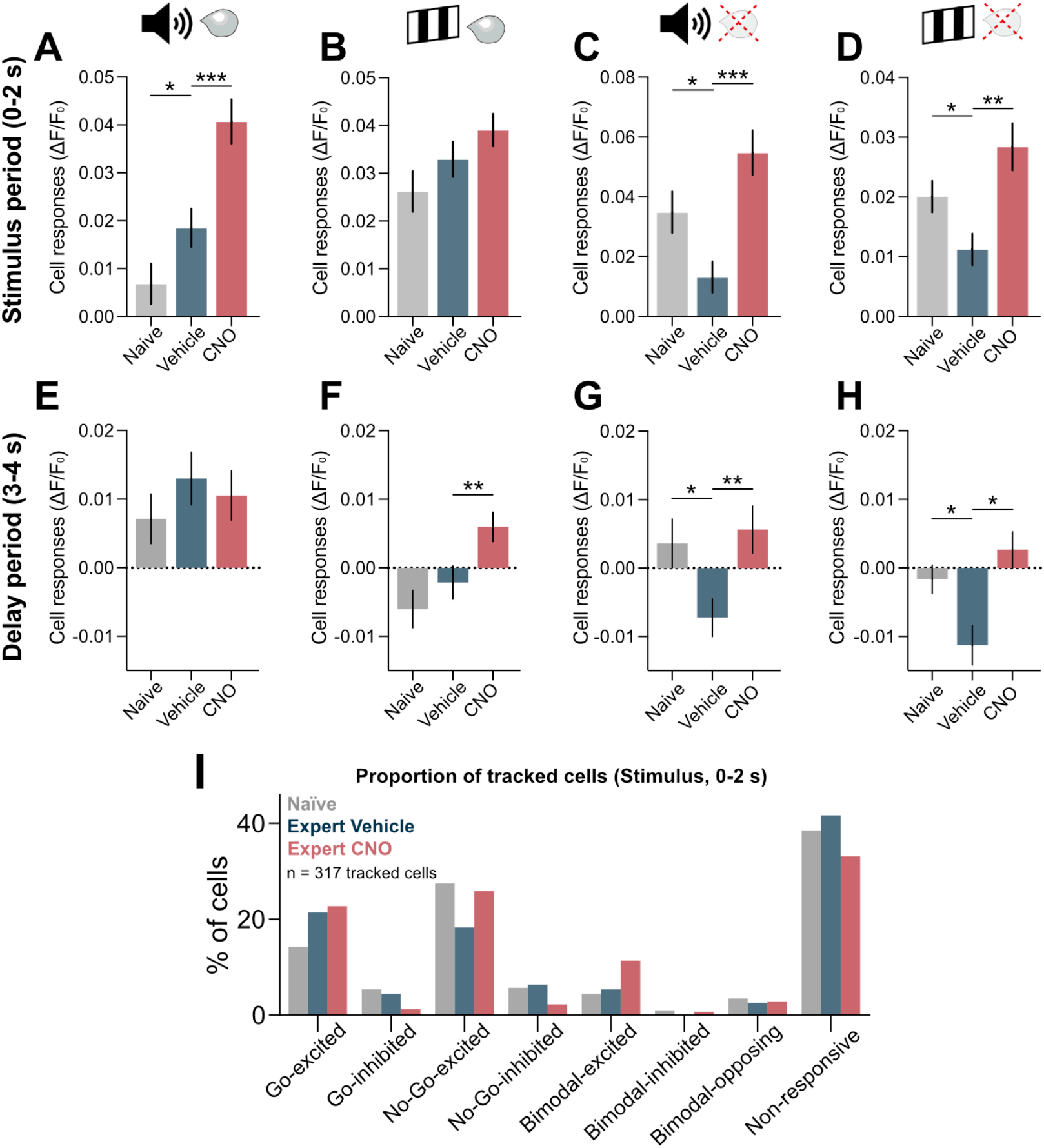
Learning- and TRN-dependent modulation of Go and No-Go cue responses. (**A-D**) Stimulus period (0-2 s). (**A**) Sound-trained mice (Go = sound): Go responses increased from naïve to expert and further under TRN blockade (Kruskal-Wallis with Benjamini-Krieger-Yekutieli correction, naïve vs expert p = 0.0402, expert vs CNO p < 0.001; n = 178 naïve, n = 188 expert cells; 5 mice). (**B**) Visual-trained mice (Go = visual): Go responses showed no significant change (n = 199 naïve cells, n = 209 expert cells; 4 mice). (**C**) Visual-trained mice (No-Go = sound): learning suppressed responses, reversed by TRN blockade (naïve vs expert p = 0.0087, expert vs CNO p < 0.001; n = 199 naïve, n = 209 expert cells; 4 mice). (**D**) Sound-trained mice (No-Go = visual): learning suppressed responses, reversed by TRN blockade (naïve vs expert p = 0.0104, expert vs CNO p = 0.0065; n = 178 naïve, 188 expert cells; 5 mice). (**E-H**) Same analysis for the delay period (3-4 s). Significant effects: **F:** Visual-trained (CNO vs expert p = 0.0057); **G**: No-Go sound (naïve vs expert p = 0.0201, CNO vs expert p = 0.0012); **H**: No-Go visual (naïve vs expert p = 0.0344, CNO vs expert p = 0.0115) (Kruskal-Wallis with Benjamini-Krieger-Yekutieli correction; n = 178 sound-trained cells for panels **A, D, E, H** and n = 209 visual-trained cells for panels **B, C, F, G**). (**I**) Proportion of responsive cells across naïve, expert vehicle and expert CNO sessions (n = 317 cells from 9 mice). Data show mean ± SEM. Statistical significance: *p<0.05, **p<0.01, ***p<0.001.

**Suppl Figure 9.**
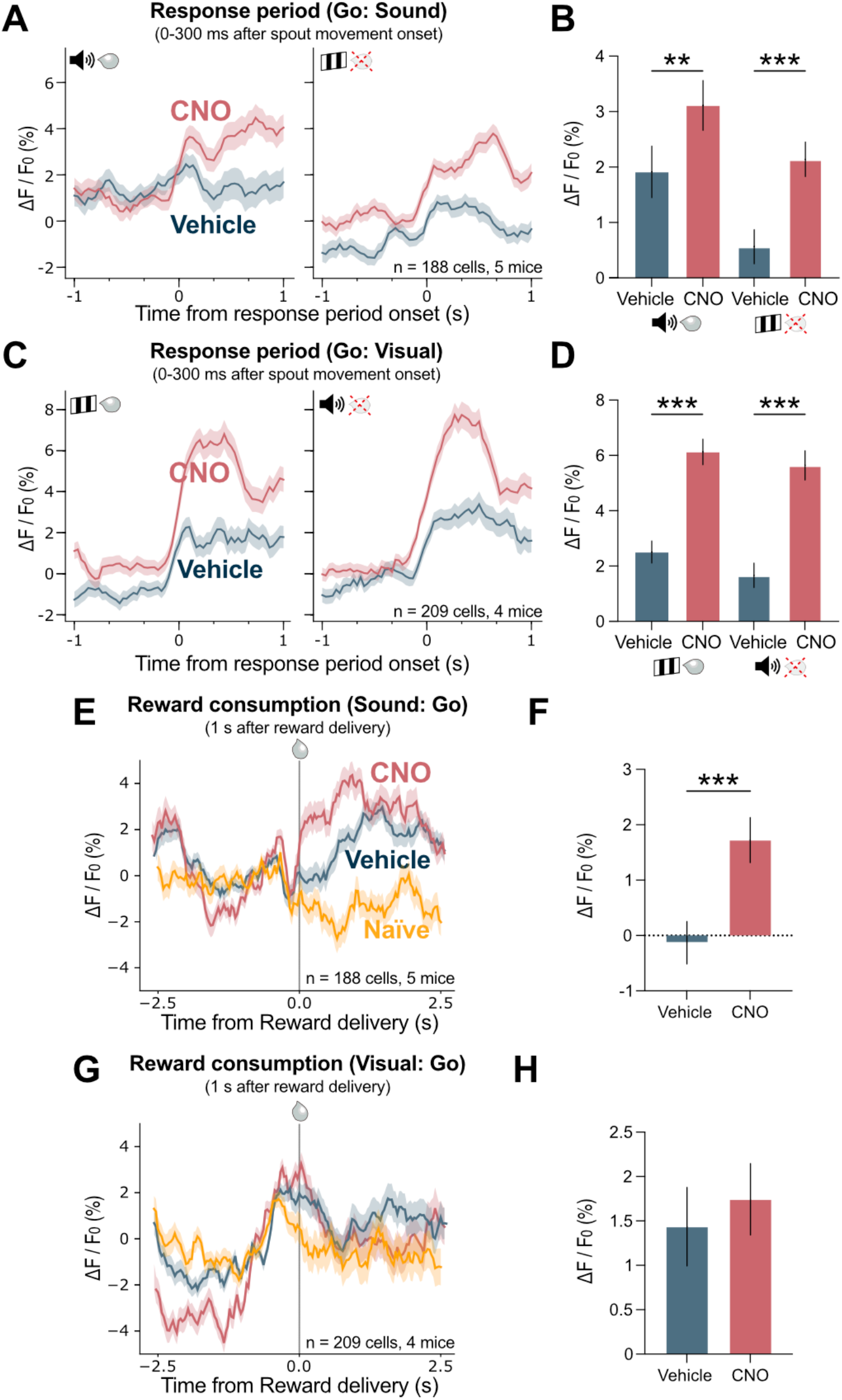
TRN modulates MGB during response period onset and reward consumption. (**A**) Mean MGB activity at response period onset in sound-trained mice (Go: sound) under vehicle and CNO conditions, shown separately for Go (left) and No-Go (right) trials (n = 188 cells from 5 mice). (**B**) TRN blockade enhanced MGB responses during response onset (0-300 ms after response period start) for both Go-sound and No-Go-visual trials (Mann Whitney U test; Go-sound p = 0.013, No-Go-visual p < 0.001; n = 188 cells from 5 mice). (**C**) Same analysis as in (**A**) for visual-trained mice (Go: visual, n = 209 cells from 4 mice). (**D**) TRN blockade enhanced response-onset activity for both Go-visual and No-Go-sound trials (Mann Whitney U test; both p < 0.001; n = 209 cells from 4 mice). (**E**) Mean MGB responses aligned to reward delivery in sound-trained mice (Go: sound) under vehicle and CNO conditions (n = 188 cells from 5 mice). (**F**) Reward consumption responses were significantly larger under TRN blockade compared to vehicle (Wilcoxon matched pairs test, p < 0.001, 188 cells from 5 mice). (**G**) Mean MGB responses aligned to reward delivery in visual-trained mice (Go: visual) under vehicle and CNO conditions (n = 209 cells from 4 mice). (**H**) Reward consumption responses in visual-trained mice under vehicle and CNO conditions (209 cells from 4 mice). Data show mean ± SEM. Statistical significance: *p<0.05, **p<0.01, ***p<0.001.

